## Supplementary Figures and Tables for "LSD1 acts as an epigenetic barrier against glucocorticoid-induced atrophy and exercise-induced hypertrophy in skeletal muscle"

### Supplemental Figure Legends

**Supplemental Figure 1. Generation of LSD1-mKO mice.**

(**A**) LSD1-mKO mice. Upon tamoxifen administration, LSD1 was depleted in differentiating and differentiated muscle cells, in which human ACTA1 promoter was activated. We performed the Dex administration and VWR tests, which induced muscle-site dependent adaptive responses. LSD1-mKO mice were generated by crossing *Lsd1*-floxed mice and knock-in mice expressing a CreERT2 recombinase under the control of human *ACTA1* gene promoter. (**B**) LSD1 mRNA expression in WT (n = 4) and LSD1-mKO (n = 4) mice. qRT-PCR values were normalized to the *36B4* gene and presented as the fold differences, compared with those in WT samples. (**C**) LSD1 protein expression in muscles from LSD1-mKO mice. Values are mean ± SD. $^{**}p < 0.01$. ACTA1, actin alpha 1; TA, tibialis anterior muscle; Gas; gastrocnemius muscle; EDL, Extensor Digitorum Longus Muscle; Sol, soleus muscle; WAT, white adipose tissue; BAT, brown adipose tissue.

**Supplemental Figure 2. Effect of Dexamethasone dosage on atrophic phenotype.**

(**A**) Experimental design: 8-week-old male WT mice were injected with PBS or Dex (1 or 5 mg/kg) for 7 d, consecutively. To arrange an experimental setting compatible with the KO mouse tests, we administered tamoxifen intraperitoneally to WT mice for 5 consecutive days and the PBS/Dex injection started at the fourth day of tamoxifen administration. (**B, C**) Body weight change (B) from day 0 to day 6 of the experiment in figure S2a and tissue weights (C) in mice treated with PBS (n = 6), 1 mg/kg BW (n = 7) or 5 mg/kg BW (n = 5) Dex. (**D**) Changes in body weight during the Dex administration period (day 3-9) in WT (n = 40) and LSD1-mKO (n = 44) mice. (**E**) Liver weight in WT (n = 8) and LSD1-mKO (n = 9) mice without Dex administration, and WT (n = 34) and LSD1-mKO (n = 39) mice with Dex administration. (**F**) Treadmill performance of Dex-treated WT (n = 10) and LSD1-mKO (n = 12) mice. Running distance and time until exhaustion are shown. Values are mean ± SD. *$p < 0.05$, **$p < 0.01$.

**Supplemental Figure 3. Effects of LSD1 mKO on muscle atrophy- and hypertrophy-associated signaling pathways.**

(**A**) Western blot analysis of an autophagy protein LC3 in EDL muscles from Dex treated-WT and LSD1-mKO mice. (**B**) Autophagic signaling was evaluated based on the LC3-II/LC3-I ratio. (**C**) Western blot analysis of phosphorylated (p)-Akt, total Akt and GAPDH protein in WT and LSD1-mKO EDL muscles after Dex treatment. (**D**) Hypertrophic signaling was evaluated based on the p-Akt/total Akt ratio. Band densities were quantified by densitometry. Values are mean ± SD, and shown as the fold difference against WT. (E) All-limb grip strength of WT (n = 3) and LSD1-mKO (n = 4) mice.

**Supplemental Figure 4. Increase in the number of slow fibers in Dex-treated LSD1-mKO mice.**

(**A**) Single color images of type I and IIA fibers shown in **Fig. 1D**. Scale bars, 300 μm. (**B**) (upper panel) The cross-sectional areas and numbers of each fiber type were determined by analyzing the individual fibers that were visualized by laminin staining. (lower panel) Fiber areas depicted in blue (type I), yellow (IIA), and magenta (IIB+IIX) were detected by a BZ-X Analyzer software (KEYENCE). (**C**) Staining of individual fiber types in Gas muscles from WT and LSD1-mKO mice after tamoxifen administration (without Dex treatment). Scale bars, 100 μm. Representative images are shown. (**D**)

Staining of the nuclei in Gas muscles from Dex-treated WT and LSD1-mKO mice. Note that DAPI-positive nuclei are located at the periphery of the fibers. Scale bars, 200 μm. (**E, F**) Size distributions of type I, type IIA (E), and type IIB + IIX fibers (F) (WT, n = 6; mKO, n = 6). Large type I and IIA fibers preferentially increased in LSD1-mKO muscle (E), while large type IIB + IIX fibers decreased (F) as highlighted by red bars.

**Supplemental Figure 5. Effects of LSD1-mKO on the expression of atrophy-, hypertrophy-, and fiber type-associated genes.**

(**A, B**) Gene expression profiles in the muscles from Dex-treated WT (n = 8) and LSD1-mKO (n = 11) mice. The expression of hypertrophy-associated genes in the Gas muscle (A) and fiber type-specific genes in the TA muscle (B) are shown. qRT-PCR values are shown as the fold differences against those in the WT. (**C, D**) Gene expression profiles in the muscle from WT (n = 3) and LSD1-mKO (n = 3) mice without Dex treatment. Gas and TA muscles were dissected from the mice 4 weeks after the start of tamoxifen administration. The expression of atrophy-associated genes (C) and fiber-type specific genes (D) are shown. qRT-PCR values are shown as the fold differences against the WT. Full descriptions of gene symbols are provided in the **Supplemental Table 3**. Values are mean ± SD *$p < 0.05$, **$p < 0.01$.

**Supplemental Figure 6. Transcriptome analysis of the Sol muscle in LSD1-mKO mice after Dex treatment.**

(**A**) Comparison of the transcriptome data of the muscles from the Dex-treated WT (n = 3) and LSD1-mKO (n = 3) mice. The Magenta and cyan dots represent the genes that were significantly upregulated and downregulated, respectively, in LSD1-mKO muscles, respectively (FDR > 0.05). (**B**) Comparison of transcriptome profiles of Gas and Sol muscles from Dex-treated LSD1-mKO mice. Genes that were significantly upregulated or downregulated in the LSD1-mKO Gas muscle are aligned in the heatmap according to the -logFC values against those in the WT. Alongside the Gas heatmap, -logFC values of corresponding genes in the Sol (mKO *vs*. WT) are indicated. (**C, D**) The expression of the atrophy-associated genes (C) and fiber type-specific genes (D) in the Sol muscle from Dex-treated WT (n = 8) and LSD1-mKO (n = 11) mice. qRT-PCR values are shown as the fold differences against those in the WT. Full descriptions of gene symbols are provided in the **Supplemental Table 3**. Values are mean ± SD. $^*p < 0.05$, $^{**}p < 0.01$.

**Supplemental Figure 7. Foxk1 cooperates with LSD1 to control the expression of atrophy-associated genes.**

(**A**) Enrichment of LSD1 at the atrophy gene loci in C2C12 myoblasts. ChIP-seq data reported by Tosic M *et al.* (Nat. Commun. 9, No. 366, 2018) was analyzed by Integrative Genomics Viewer (IGV). (**B**) Co-immunoprecipitation of Sin3A and LSD1 or Foxk1. C2C12 myotubes were treated with insulin for 6 h before harvest to enhance the nuclear retention of Foxk1. Input lane contains 10 % amount of the whole-cell extract. (**C**)Foxk1 protein expression in Foxk1-KO C2C12 cells. C2C12 cells were transfected with control or Foxk1-KO Double Nickase Plasmids and subjected to puromycin selection. (**D**) shRNA-mediated knockdown (KD) of the *Foxk1* gene in C2C12 cells. C2C12 cells expressing an shRNA against the firefly luciferase gene (shGL3) were used as control. (**E**) Expression of atrophy-associated genes in *Foxk1*-KD C2C12 cells. qRT-PCR values are shown as the fold differences against shGL3 (n = 6). Values are mean ± SD. $^*p < 0.05$, $^{**}p < 0.01$.

**Supplemental Figure 8. LSD1-mKO increased the number of oxidative fibers in slow dominant muscle after VWR training.**

(**A**) LSD1 mRNA expression in Sol muscles from WT (n = 6) and LSD1-mKO (n = 6) mice after VWR training. qRT-PCR values are normalized to the *36B4* gene and shown as fold differences against WT. (**B**) Neither the training nor LSD1 genotype affected liver weight. Sedentary WT (n = 7) and LSD1-mKO (n = 4) mice, and trained WT (n = 23) and LSD1-mKO (n = 22) mice were analyzed. (**C**) Occupancy of type IIA fiber based on the cross-sectional area (WT and mKO, n = 8). (**D**) Frequency of type I fiber based on the fiber number (WT and mKO, n = 8). (**E**) All-limb grip strength in WT (n = 10) and LSD1-mKO mice (n = 9) after VWR training. (**F**) SDH staining of the Sol muscles from trained-WT and LSD1-mKO mice. Scale bar, 100 μm. (**G**) Cumulative running activity of mice in cages with free access to a vertical running wheel for 30 d (day 4-34 in **Figure 4A**) (WT: n = 10, mKO: n = 9). Values are mean ± SD. $**p < 0.01$.

**Supplemental Figure 9. Expression of *Esrrg* and its target genes remain unaffected in LSD1-mKO Gas and TA muscles after VWR.**

The expression of *Esrrg* and its target genes in the Gas and TA muscles after VWR training (WT: n = 6, mKO: n = 6). qRT-PCR values are shown as the fold differences against the WT. Full descriptions of gene symbols are provided in the **Supplemental Table 2**. Values are mean ± SD. $*p < 0.05$, $**p < 0.01$.

**Supplemental Figure 10. LSD1 expression is decreased in the aged muscles in mice and humans.**

(**A**) Muscle weight in young (2.5 months old, n = 4) and old (28 months old, n = 3) male C57BL/6J mice. (**B**) The expression of fiber type-specific genes and atrophy-associated genes in the Gas muscles in young and old mice. qRT-PCR values are shown as fold differences against the young mice. (**C**) Expression of LSD1 protein in the EDL muscles from the young and old mice. Band densities were quantified by densitometry and normalized to histone H3. Values are shown as fold differences against the young. (**D**) Decreased expression of *Lsd1* in aged muscles. Using single cell RNA-seq data (flow cytometry) published by the Tabula Muris Senis (74), *Lsd1* expression was analyzed in Myod1-positive and Pax7-positive muscle cells from young to old mice (3-24 months old). (**E**) The expression of *LSD1* in skeletal muscles of 55-80 years-old men and women. RNA-seq data by Tumasian *et al.* (75) was analyzed. Values are mean ± SD. $R^2$: coefficient of determination. *$p < 0.05$, **$p < 0.01$.

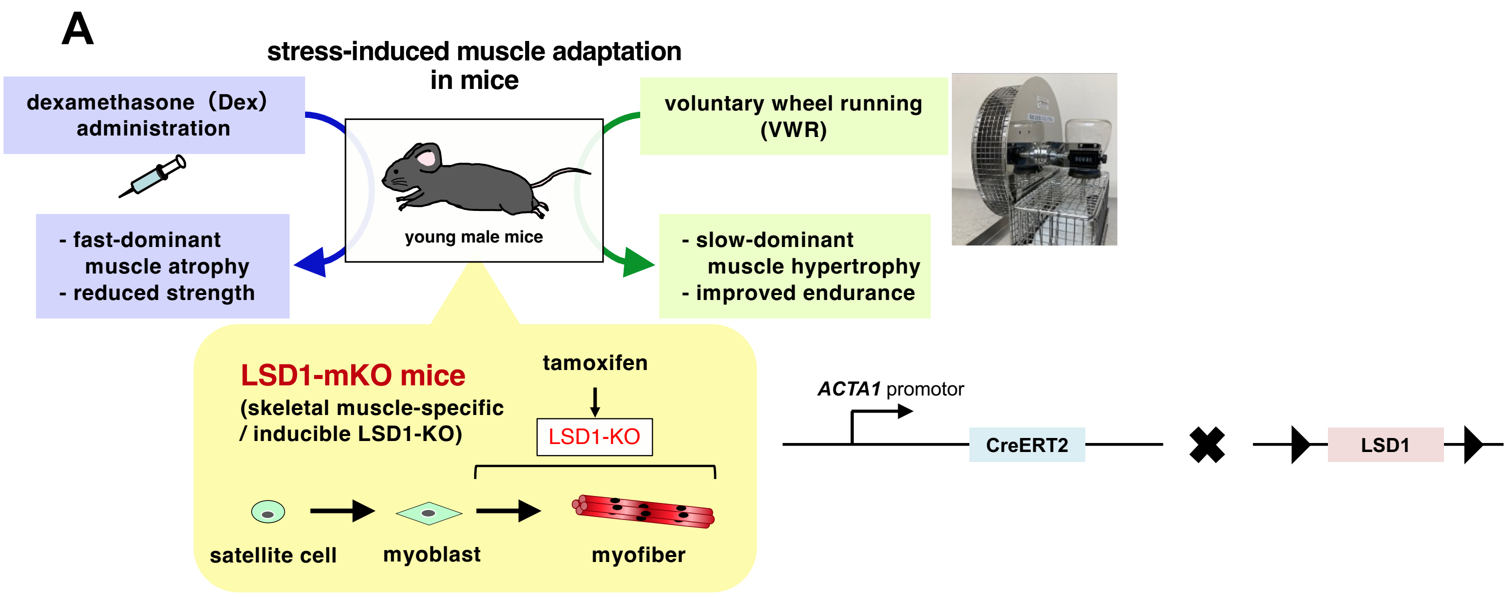


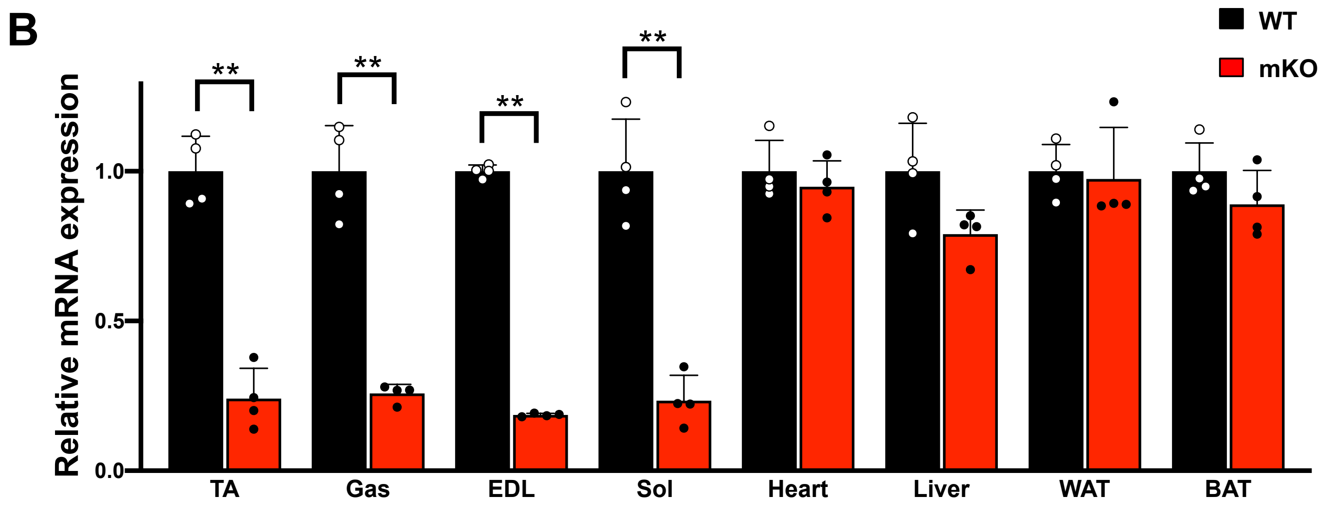


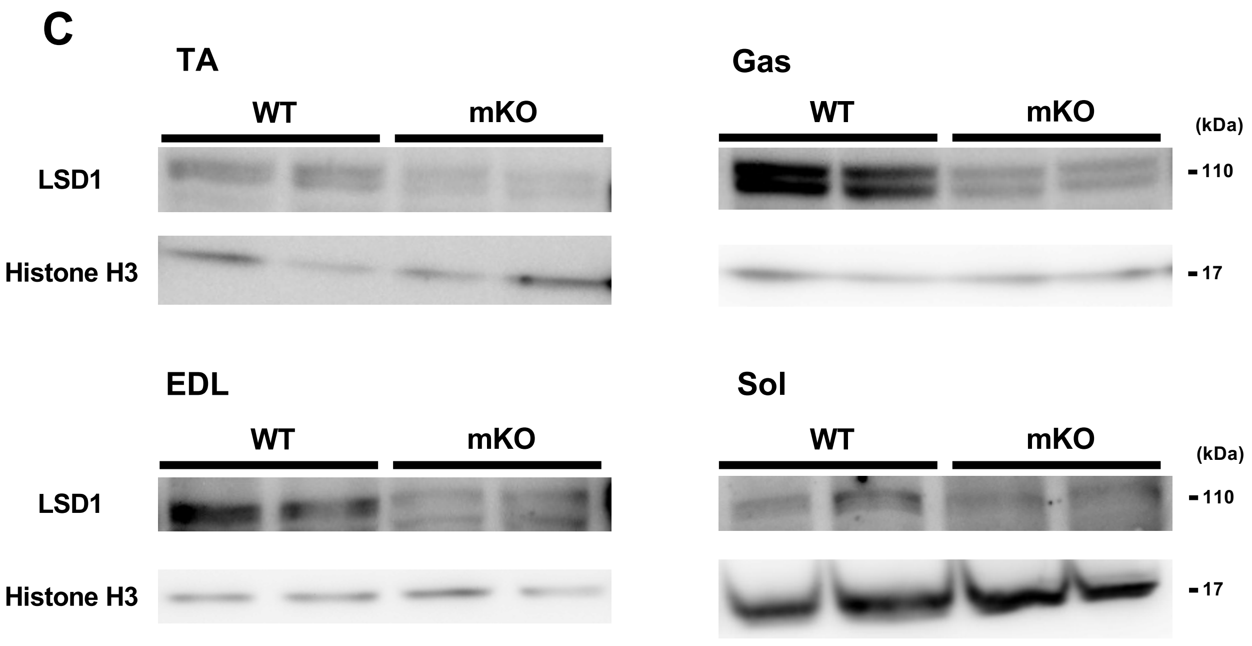


Supplemental Figure 1 (related to Fig.1)

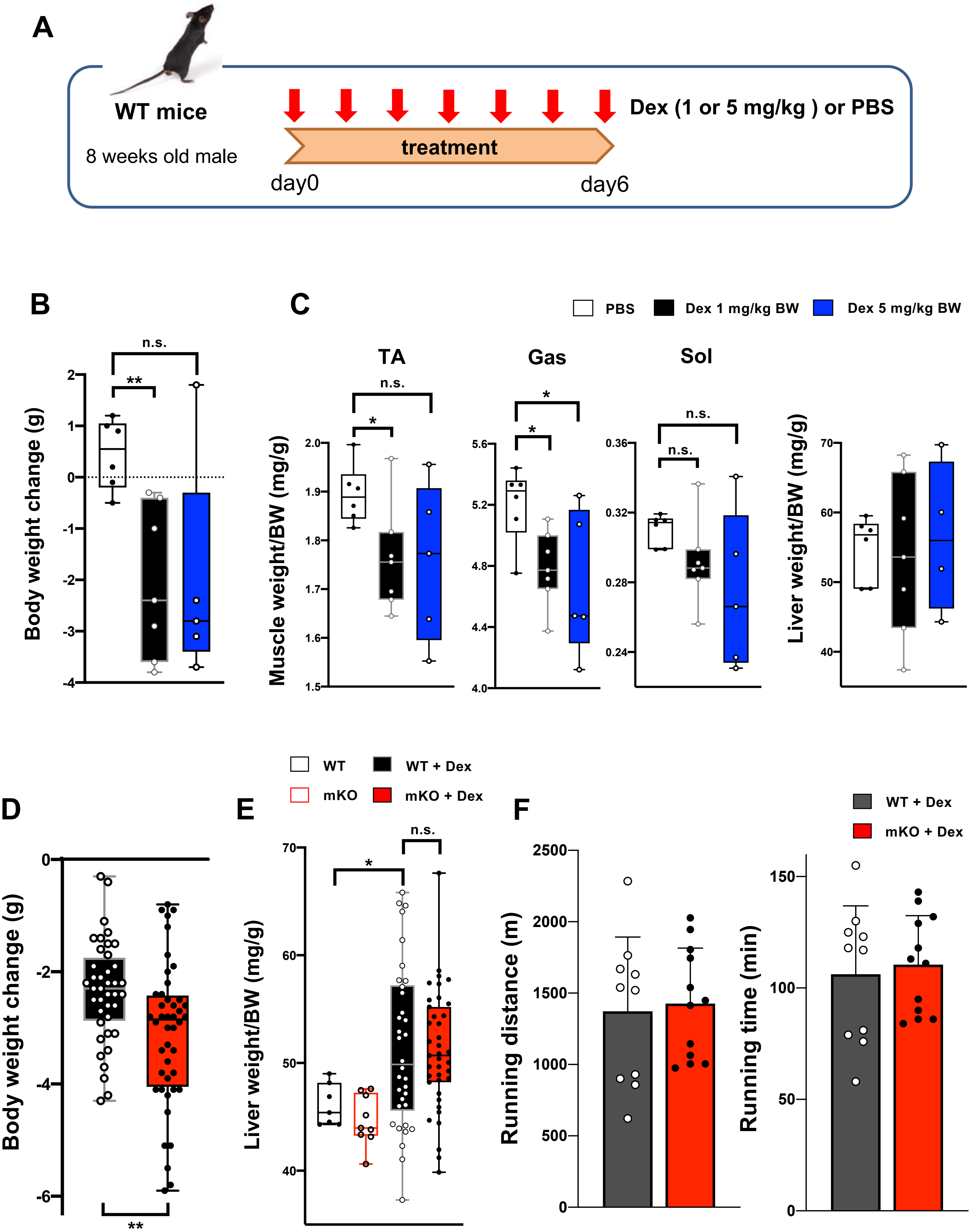


Supplemental Figure 2 (related to Fig.1)

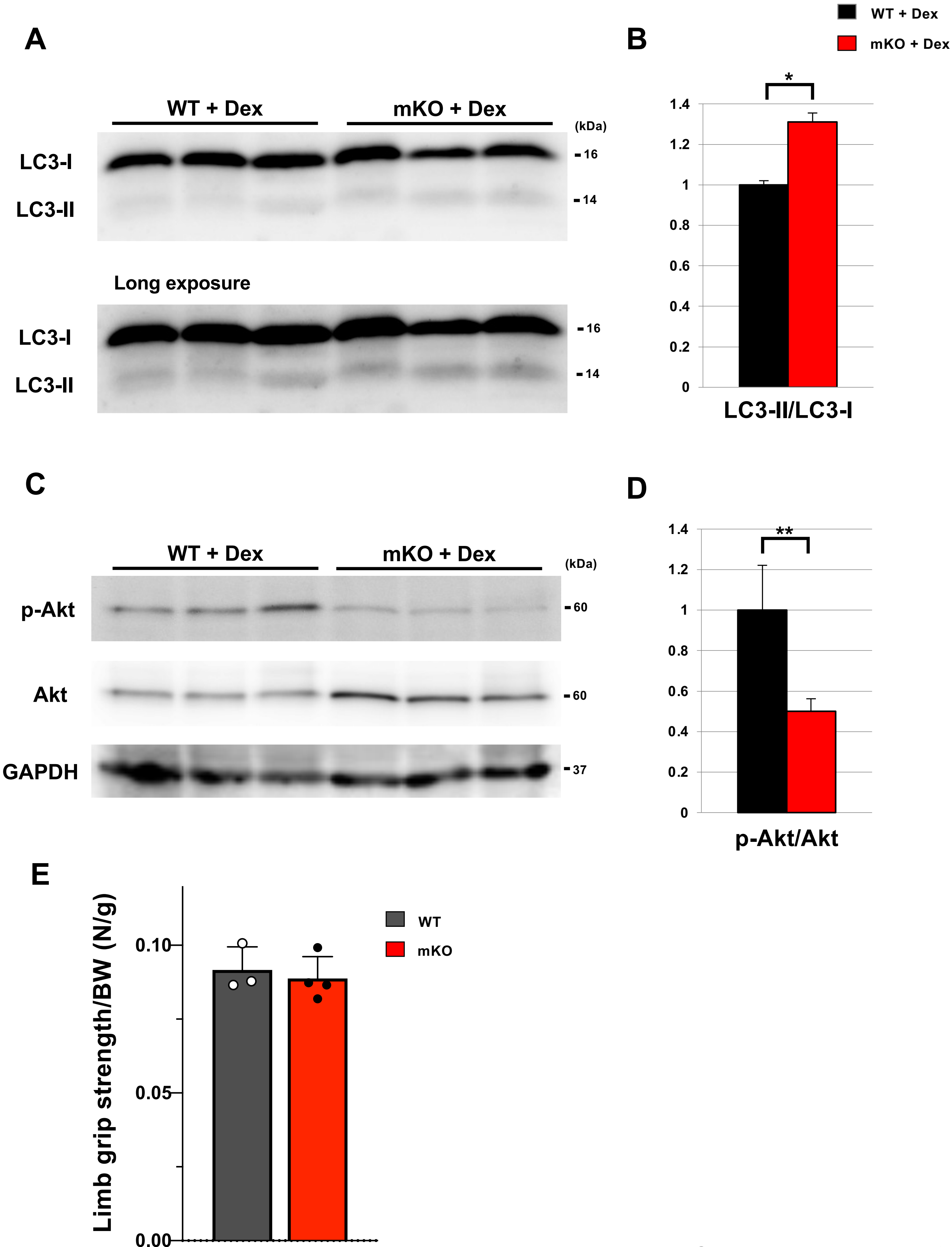


Supplemental Figure 3

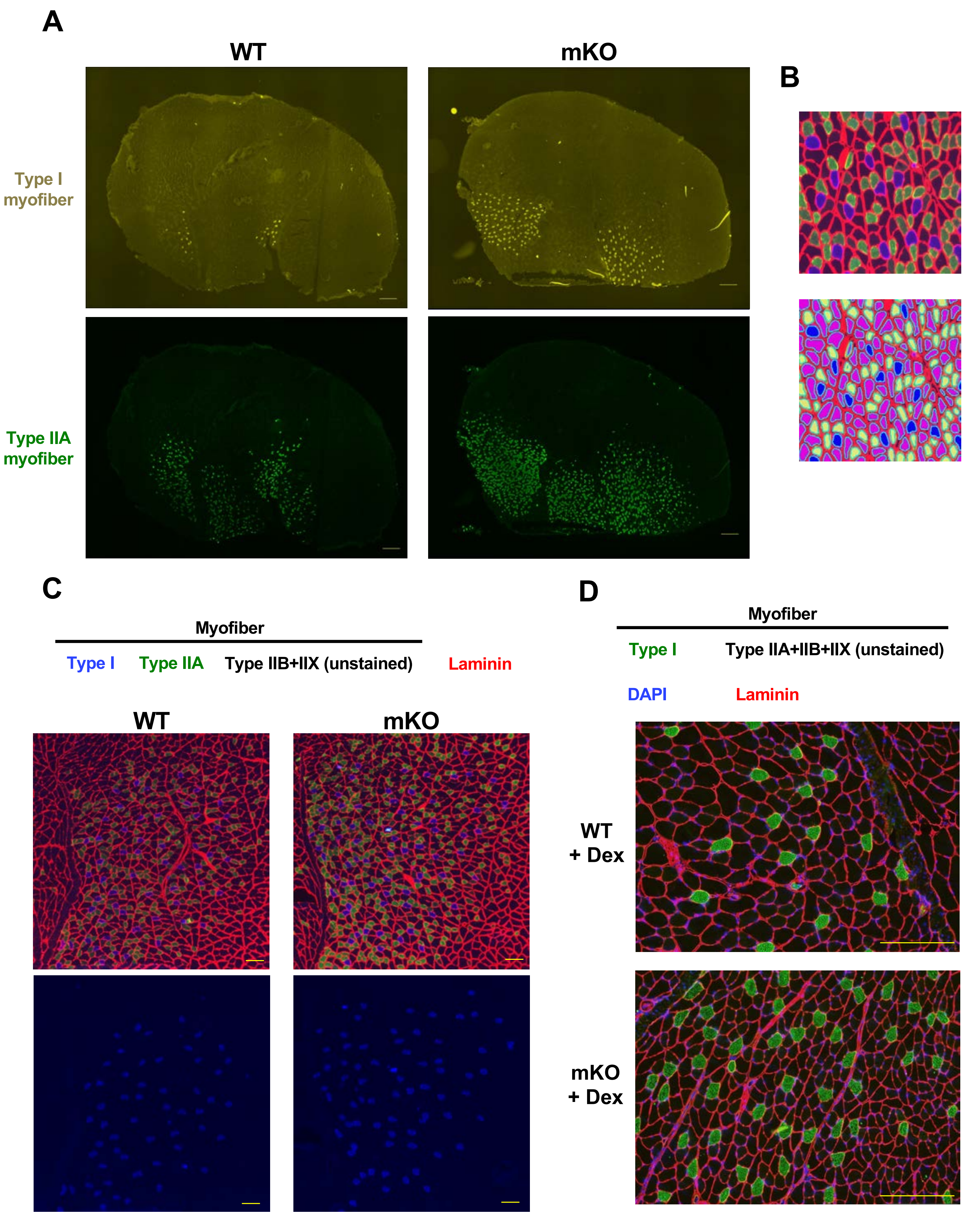


Supplemental Figure 4 (related to Fig.1)

E

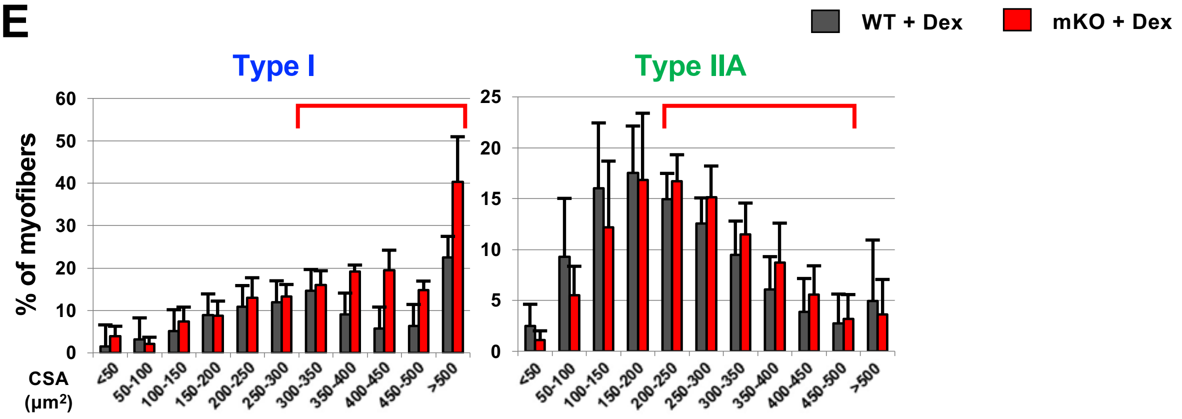


F

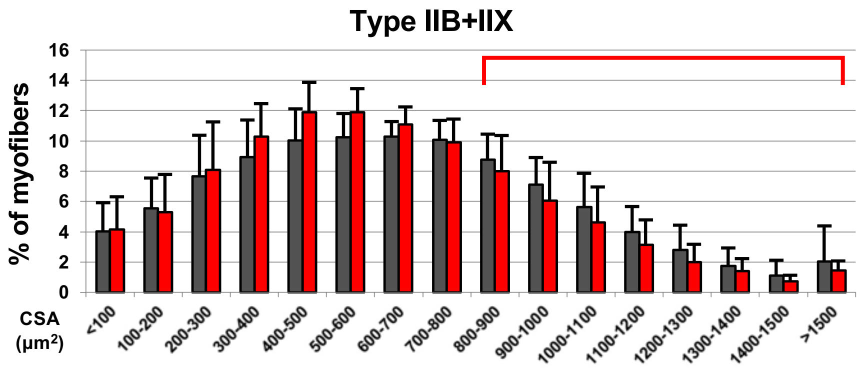


Supplemental Figure 4 (related to Fig.1)

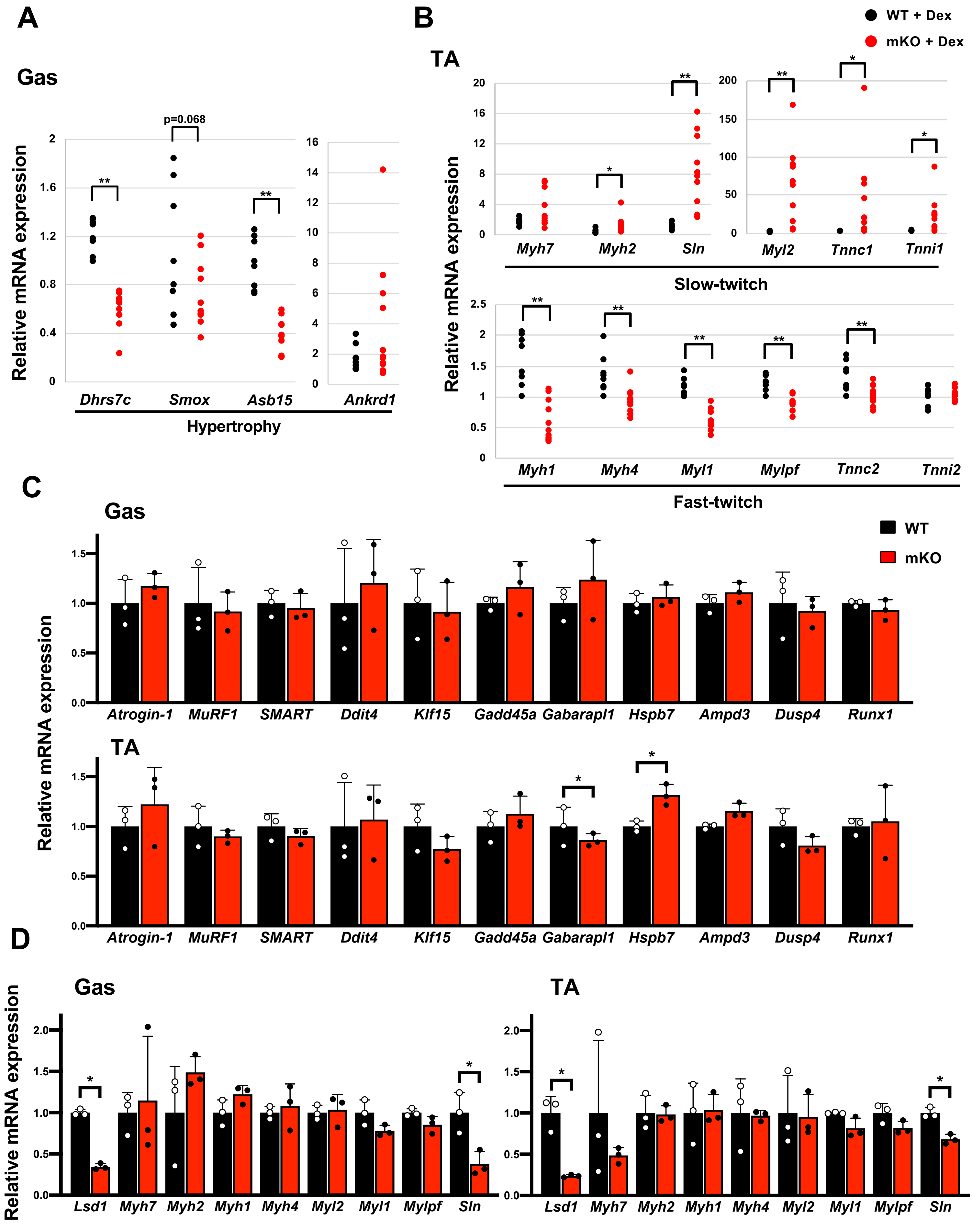


Supplemental Figure 5 (related to Fig.2)

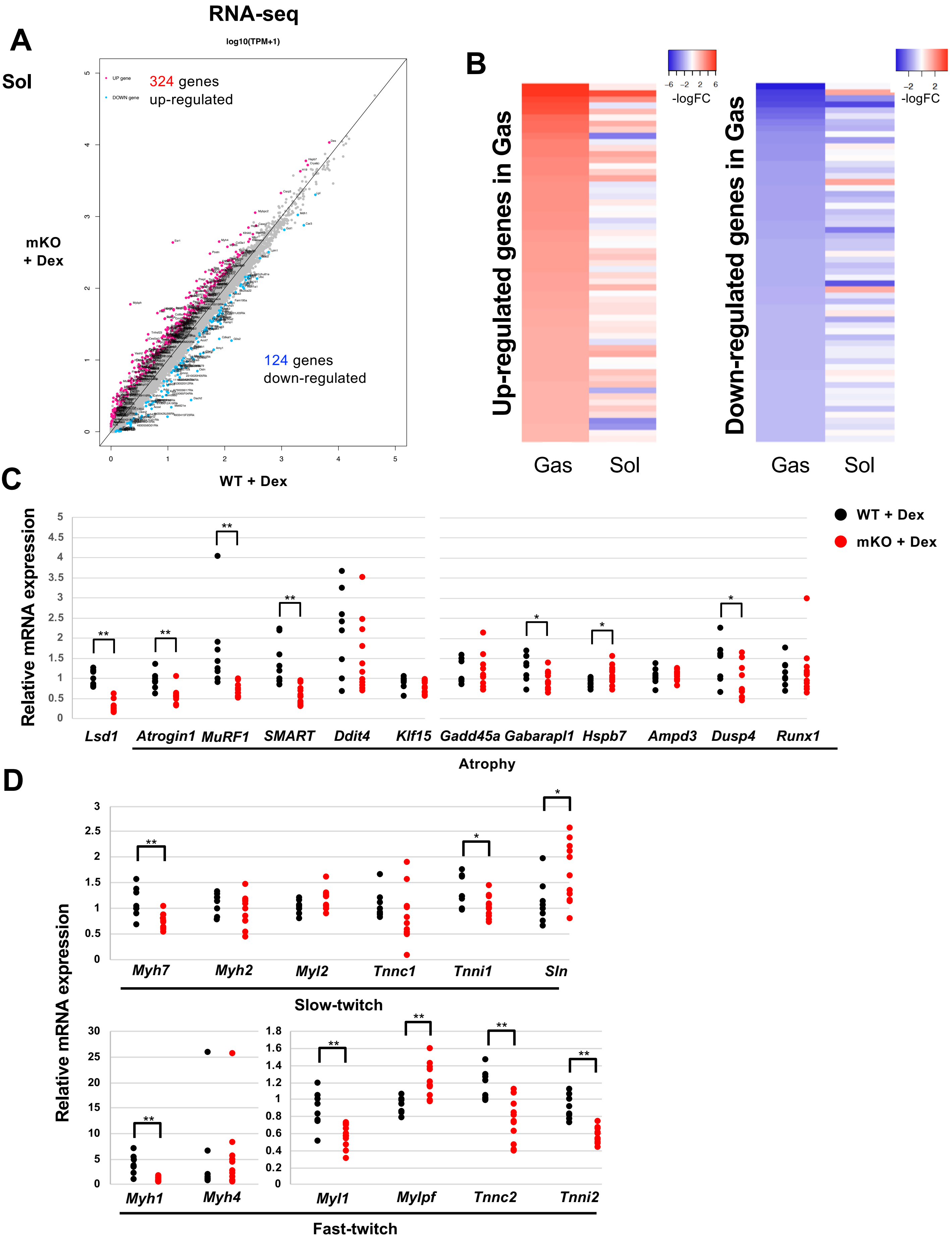


Supplemental Figure 6 (related to Fig.2)

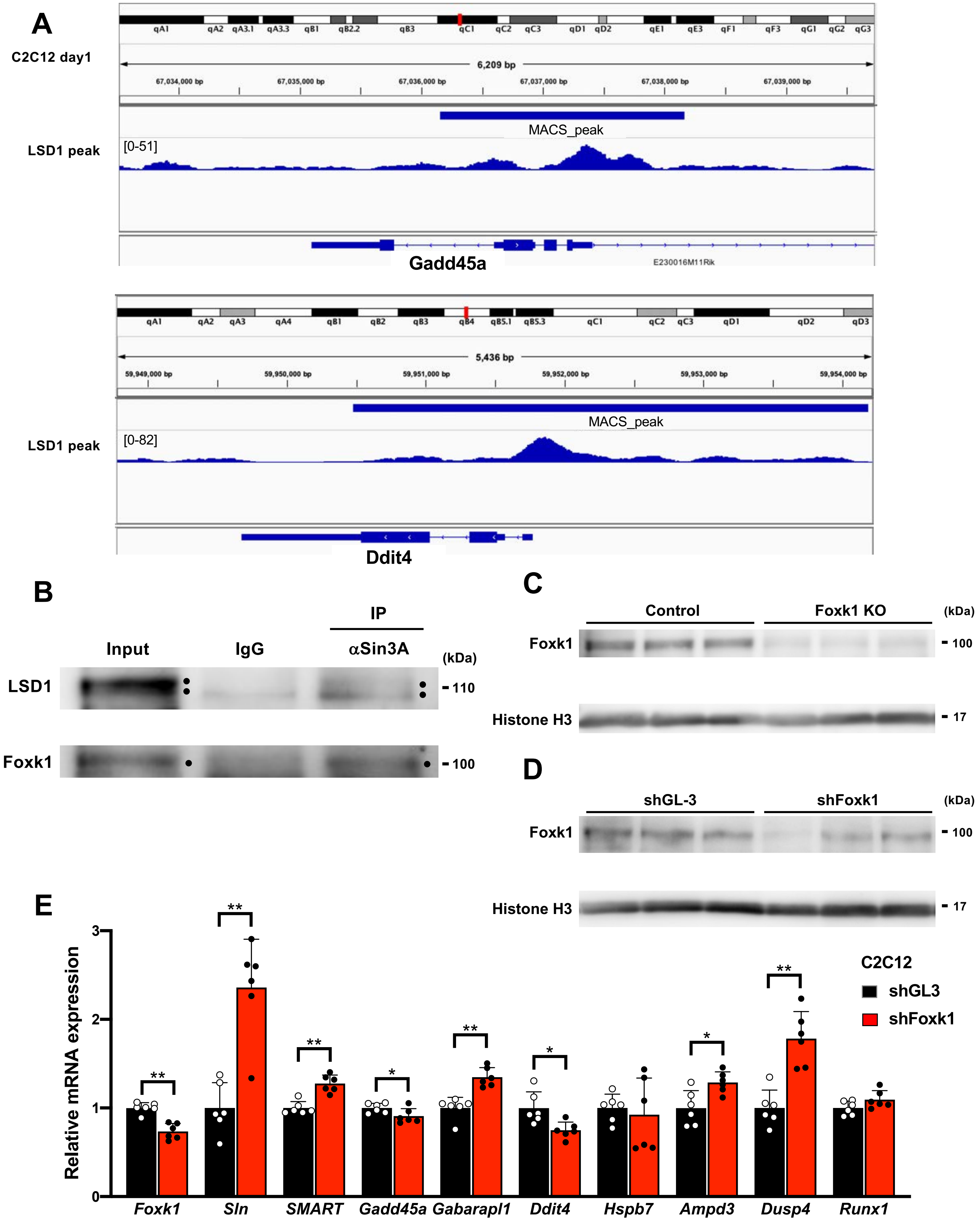


Supplemental Figure 7 (related to Fig.3)

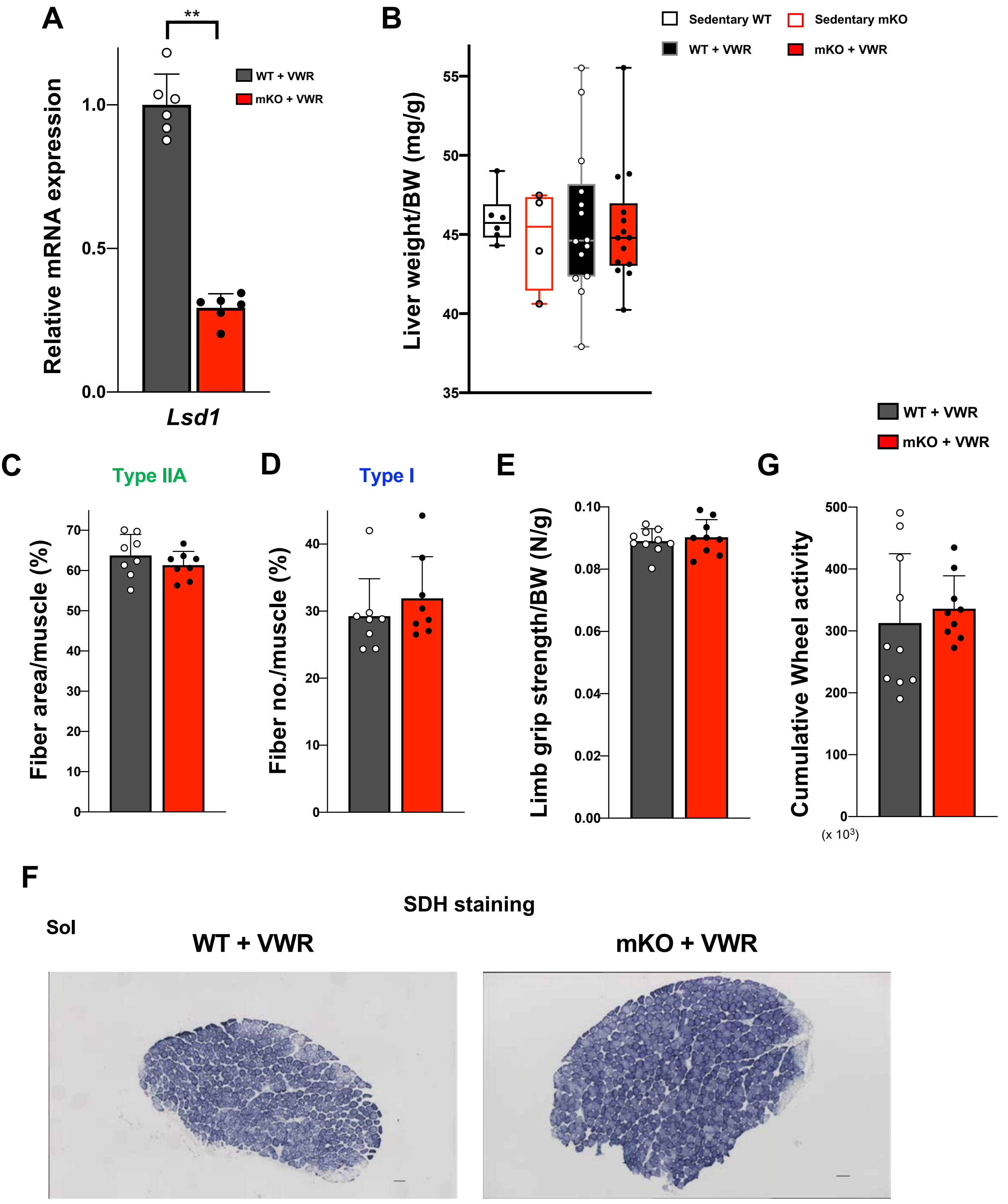


Supplemental Figure 8 (related to Fig.4)

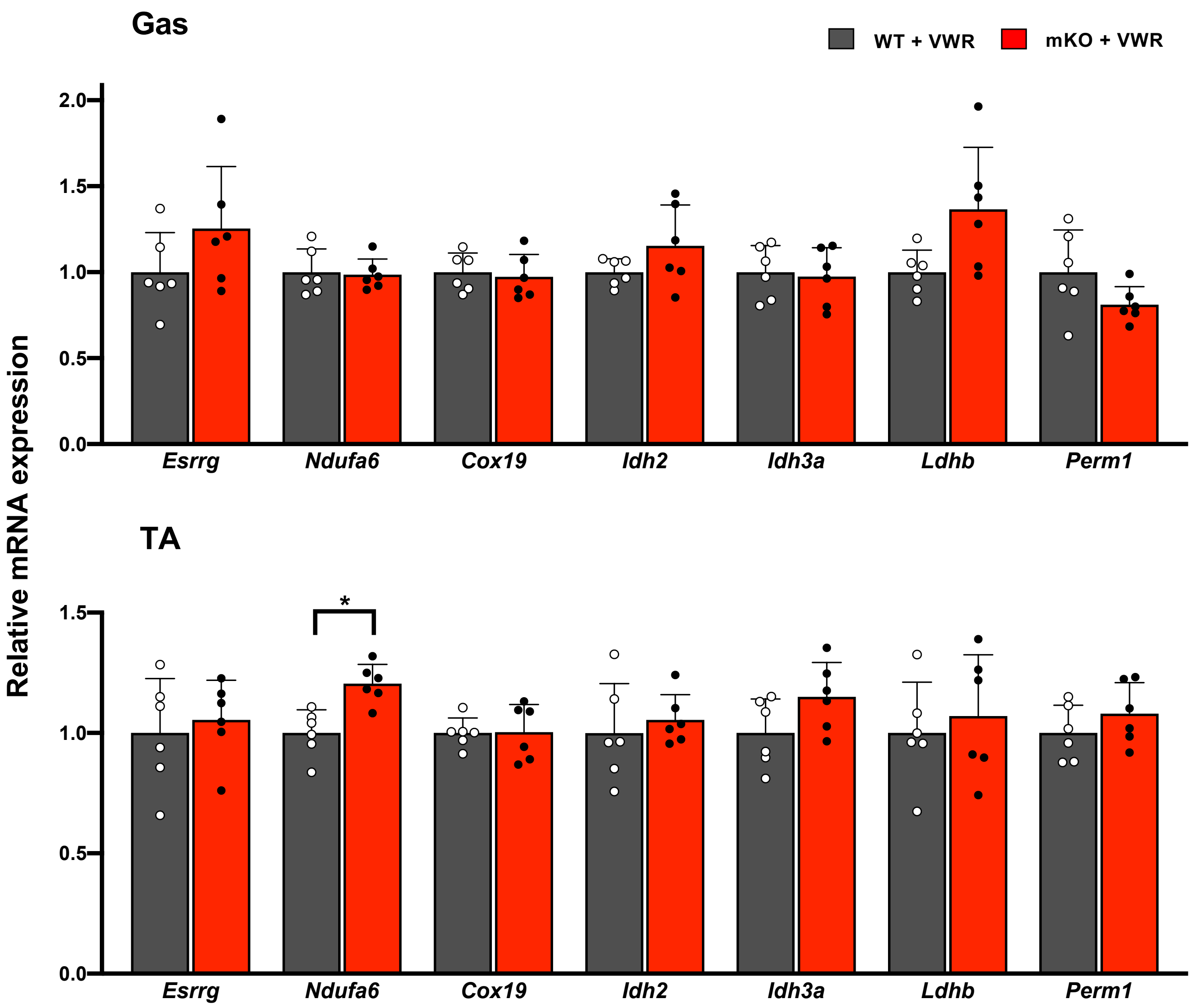


Supplemental Figure 9 (related to Fig.5)

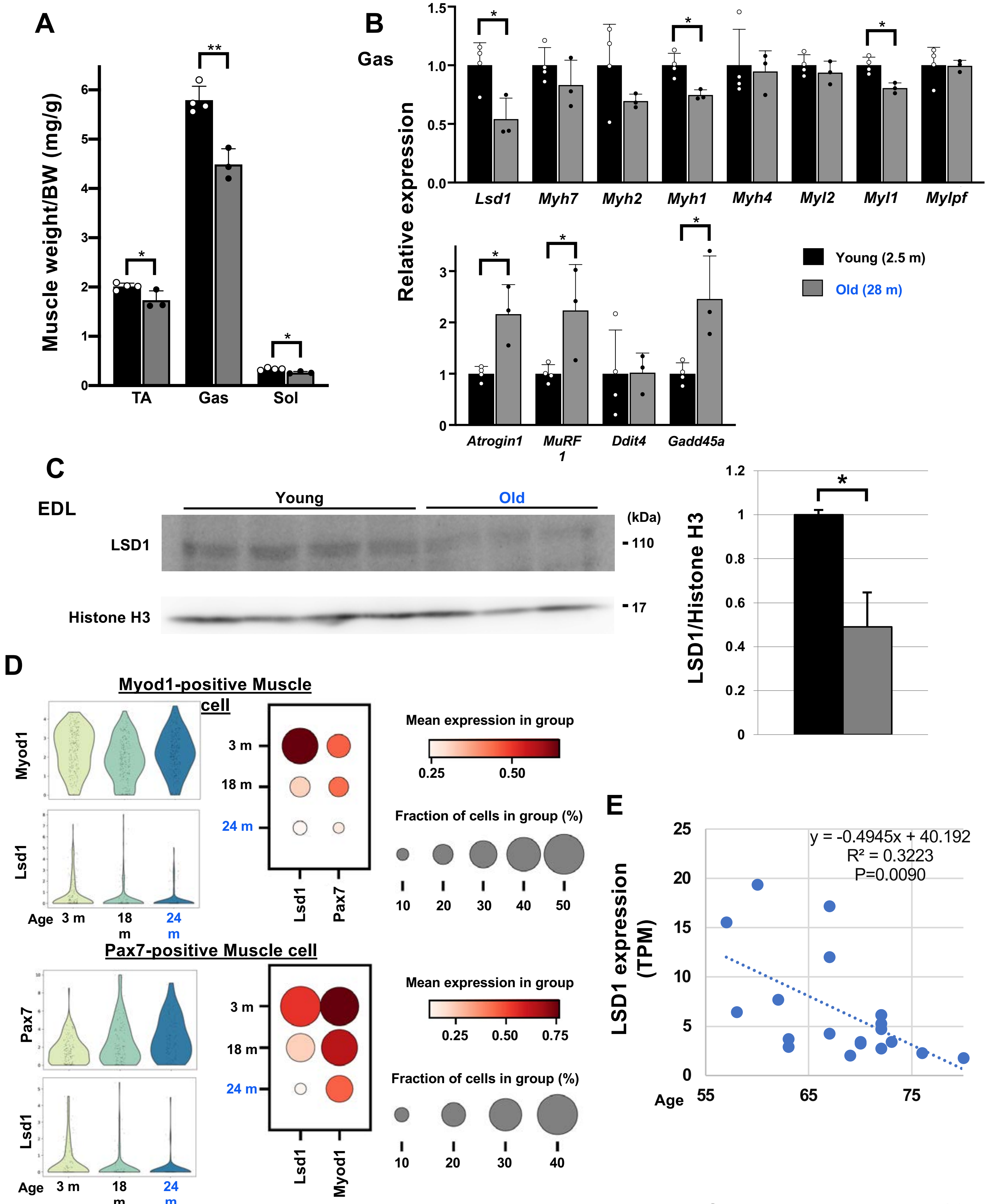


Supplemental Figure 10

Fig. 3E

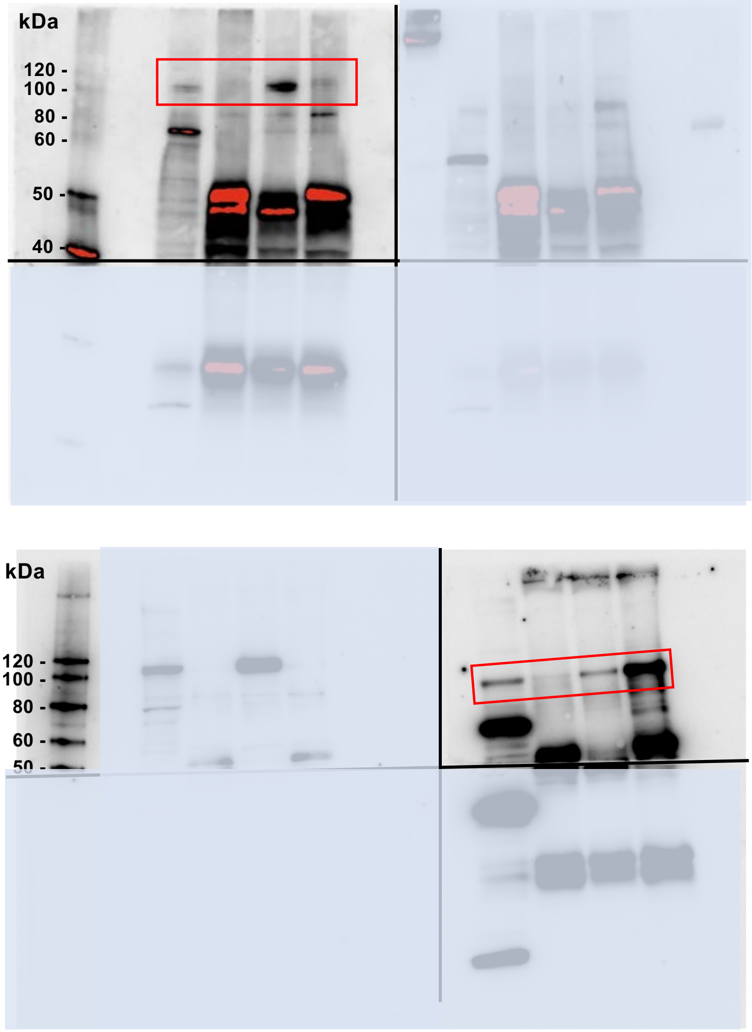


Fig. 5D

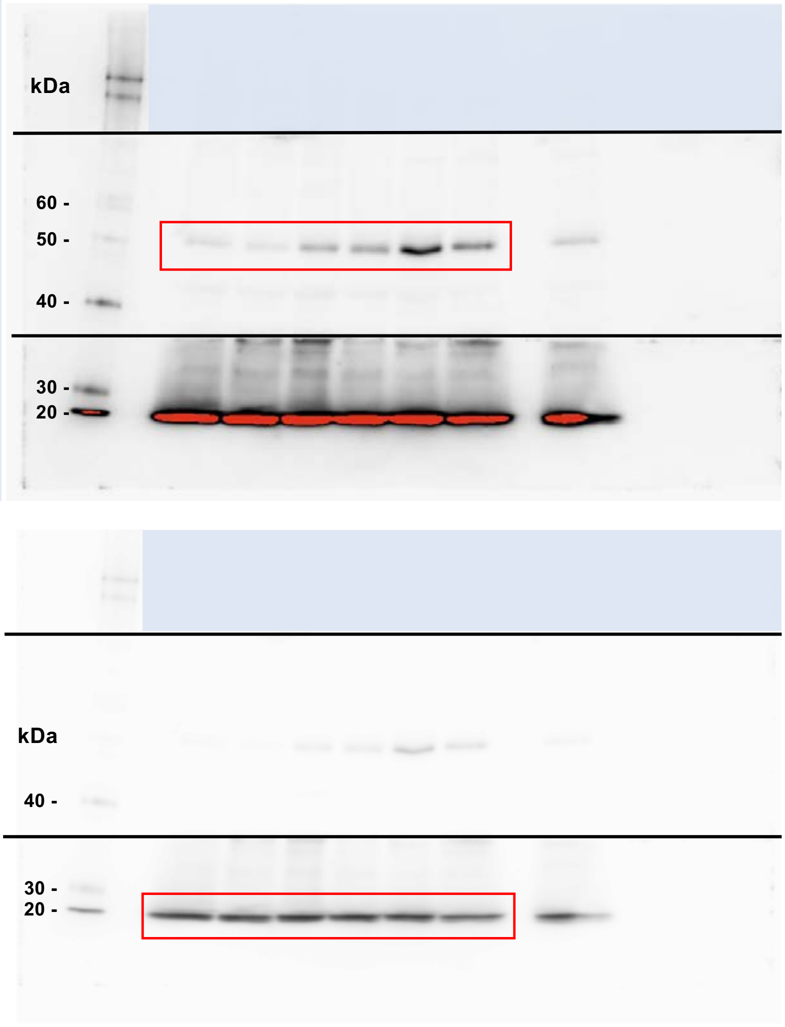


Full uncut gels

Supplemental Fig. 1C

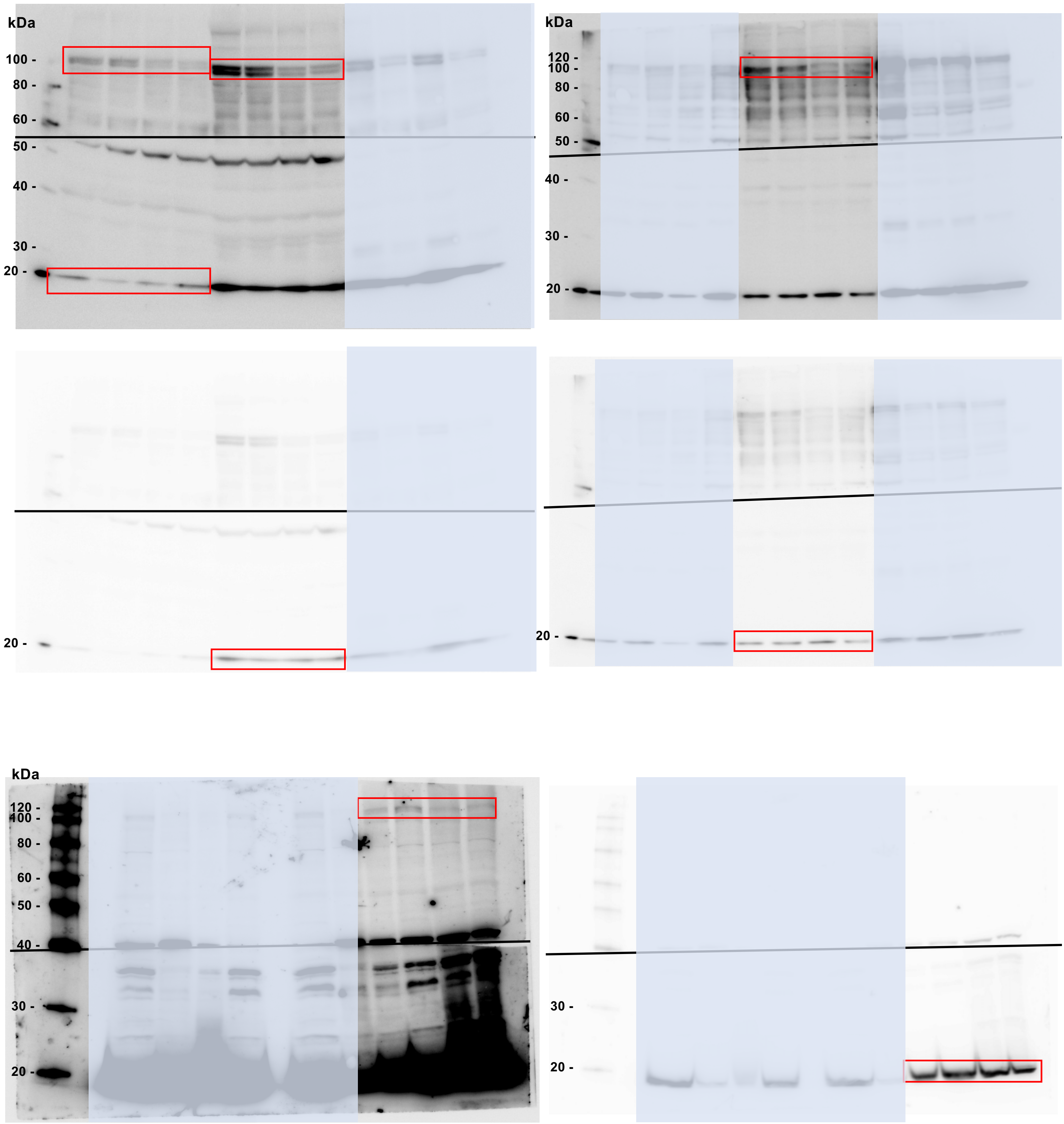


Full uncut gels

Supplemental Fig. 3A

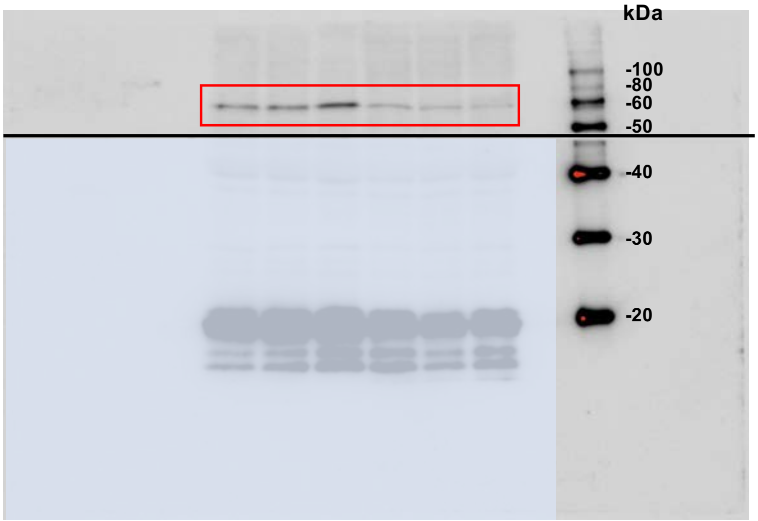


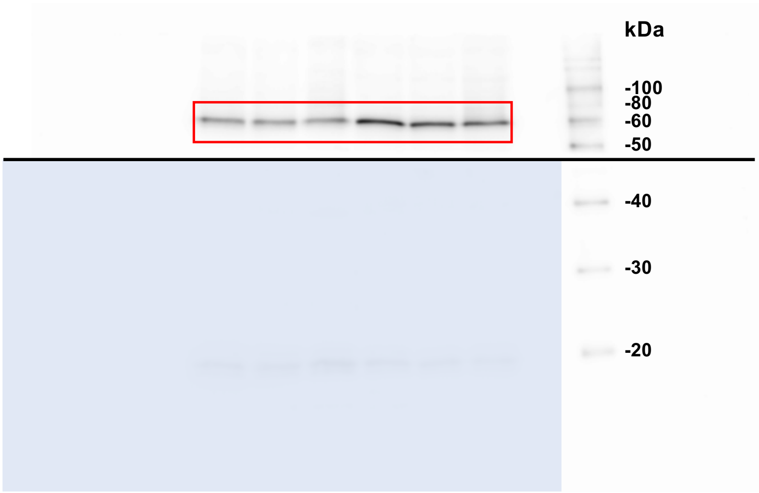


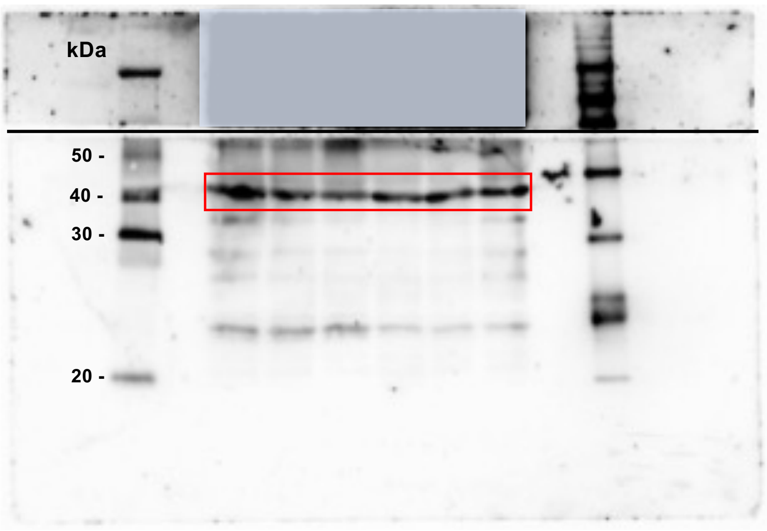


Supplemental Fig. 3C

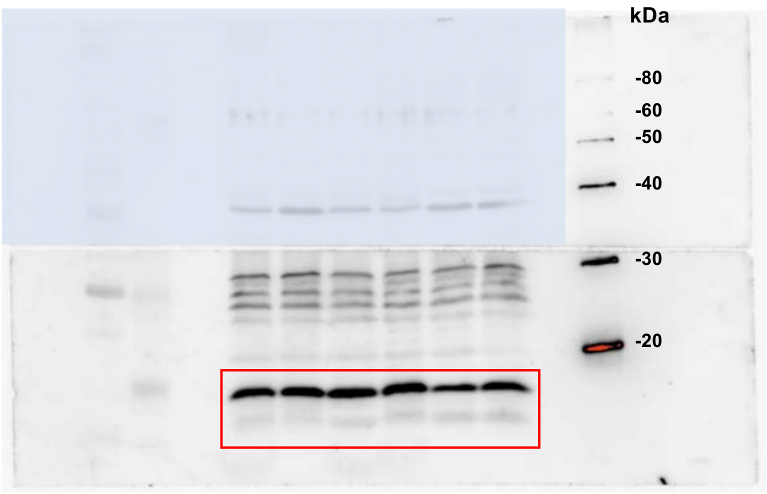


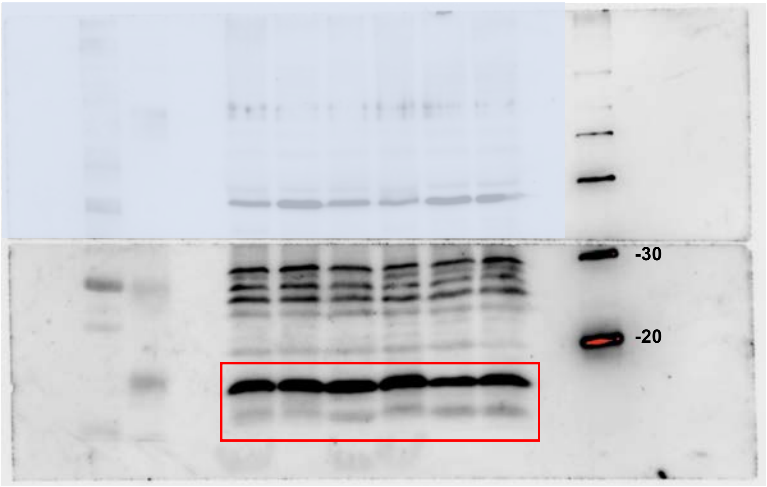


Full uncut gels

Supplemental Fig. 7B

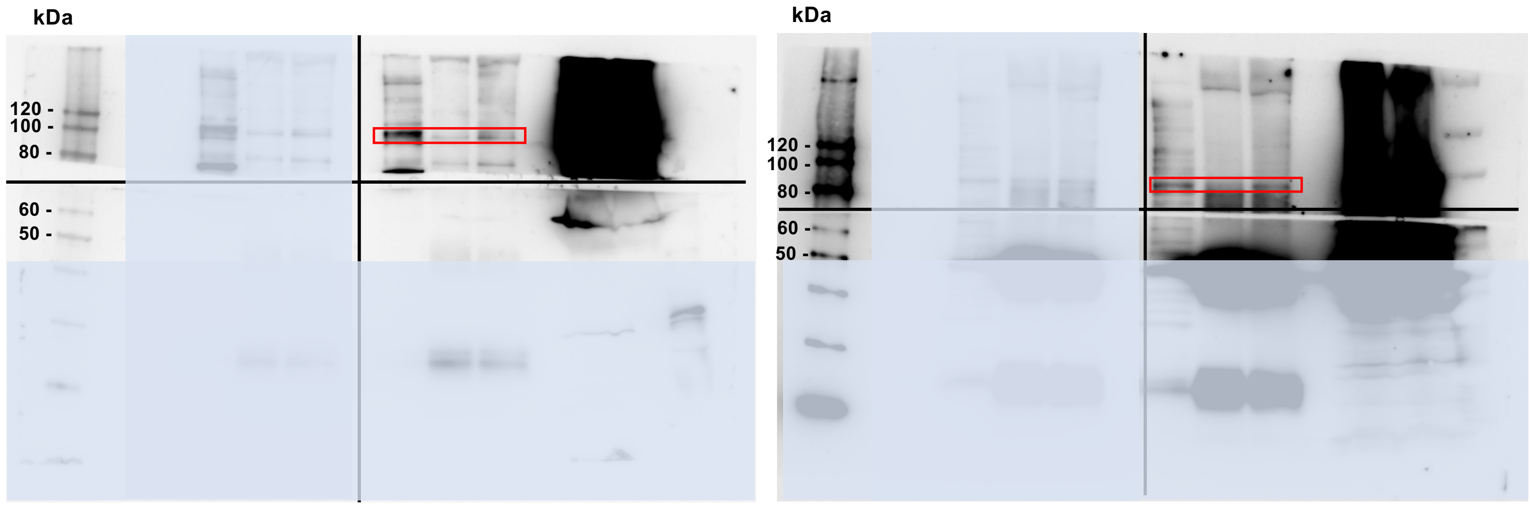


Supplemental Fig. 7C

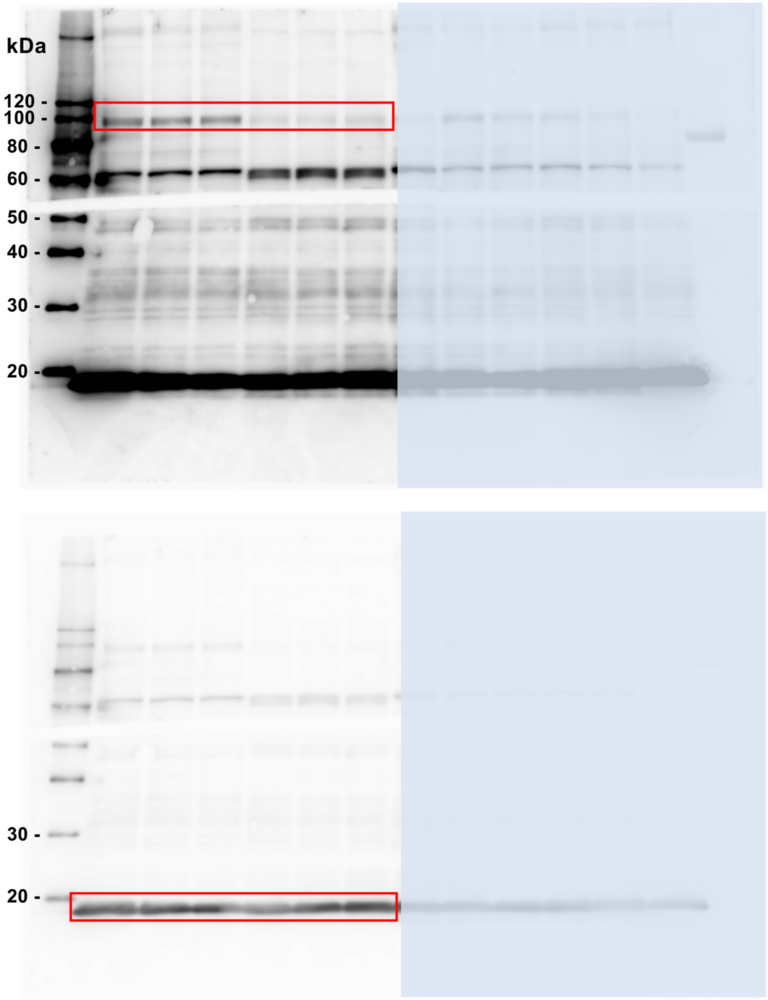


Supplemental Fig. 7D

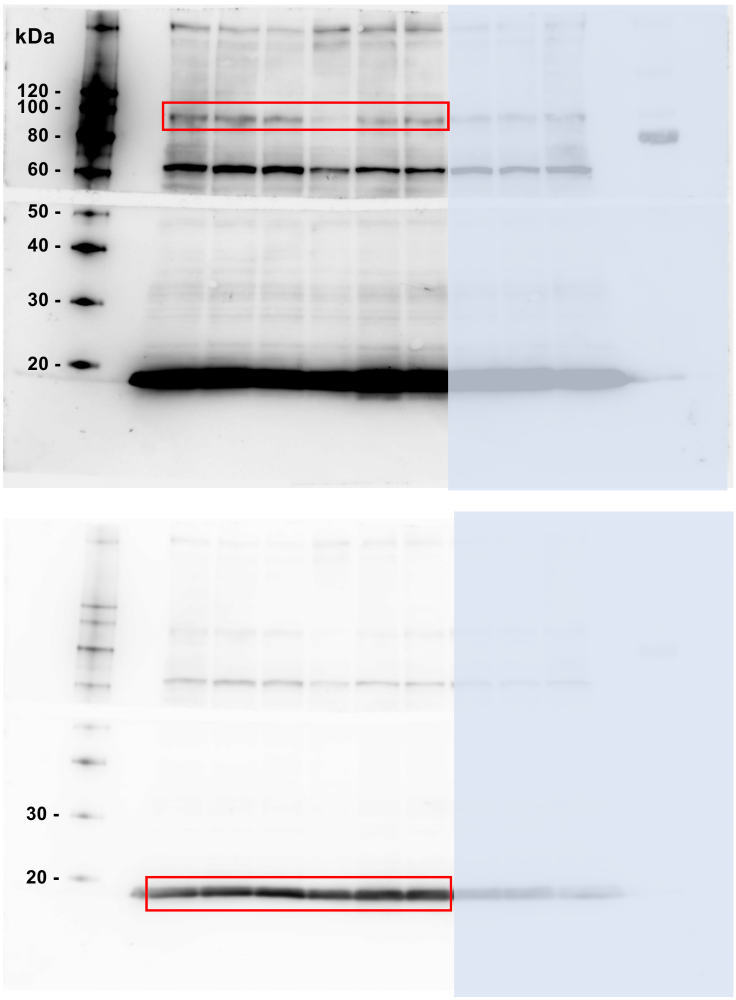


Full uncut gels

Supplemental Fig. 10C

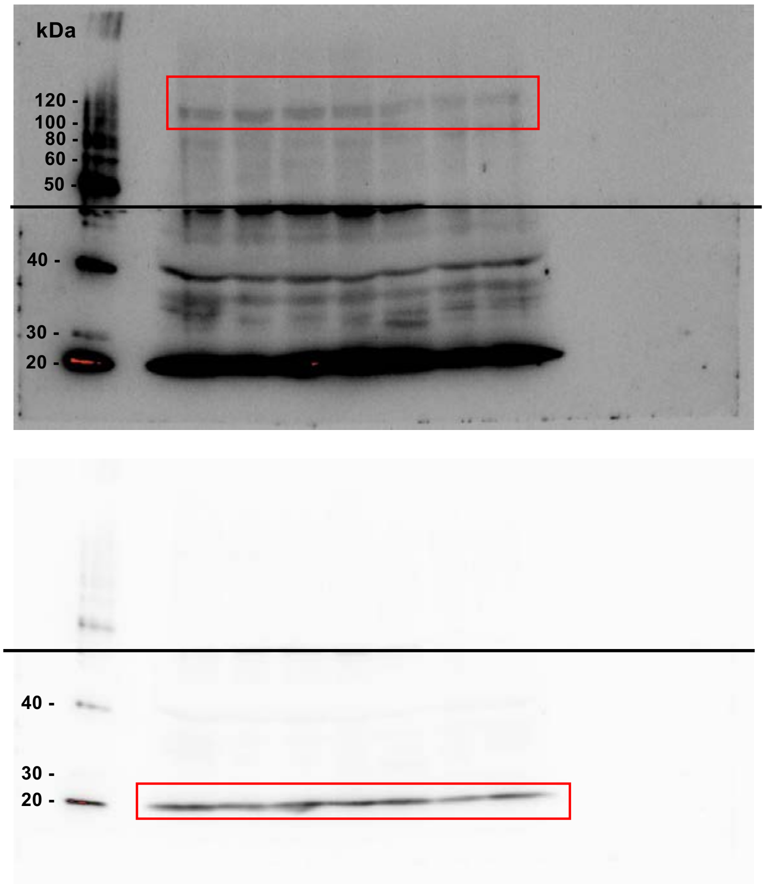


Full uncut gels

**Table S1. Top 50 differentially regulated genes in Dex-treated LSD1-mKO mice**

**Up-regulated**

| Gene Name | WT1 | WT2 | WT3 | KO1 | KO2 | KO3 | KO4 | KO5 | logFC | logCPM | PValue | FDR |
|---|---|---|---|---|---|---|---|---|---|---|---|---|
| H19 | 643.9891 | 787.649 | 529.4279 | 2913.995 | 2281.151 | 972.8129 | 892.3181 | 1137.598 | -1.32662 | 10.31067 | 9.26E-05 | 0.013782 |
| Hspb7 | 575.5697 | 588.7299 | 596.9932 | 1879.105 | 1879.718 | 1402.828 | 874.3752 | 1168.945 | -1.29542 | 10.13049 | 0.000132 | 0.018062 |
| Esr1 | 20.60644 | 15.71313 | 9.794966 | 903.0274 | 1473.462 | 650.2008 | 1132.771 | 597.4584 | -5.94525 | 9.230019 | 2.23E-42 | 5.46E-38 |
| Myl2 | 219.2646 | 146.7697 | 349.2205 | 715.7173 | 763.4889 | 342.8552 | 358.4938 | 539.0218 | -1.1904 | 8.746503 | 0.000423 | 0.042168 |
| Wnk2 | 179.8655 | 189.1269 | 52.17319 | 518.7643 | 497.9591 | 376.6834 | 284.4452 | 229.4887 | -1.44109 | 8.186018 | 2.41E-05 | 0.004682 |
| Ddit4 | 164.8515 | 63.13717 | 61.06862 | 302.7527 | 278.6113 | 795.6016 | 156.3856 | 188.7215 | -1.83661 | 7.974666 | 1.3E-07 | 6.36E-05 |
| Gabarapl1 | 134.9747 | 161.2304 | 55.97124 | 172.6182 | 211.4196 | 764.867 | 178.1539 | 185.6347 | -1.36463 | 7.865721 | 6.04E-05 | 0.009435 |
| Padi2 | 125.8051 | 103.5017 | 67.76518 | 513.8063 | 303.1889 | 250.9877 | 150.1922 | 161.0998 | -1.47746 | 7.712383 | 1.57E-05 | 0.003358 |
| Arrdc3 | 64.84228 | 29.94603 | 57.77031 | 56.31128 | 176.5353 | 457.3868 | 73.13784 | 105.3775 | -1.77274 | 6.997564 | 3.39E-07 | 0.000148 |
| Clu | 51.33977 | 51.69391 | 71.76312 | 269.9576 | 174.0247 | 91.93468 | 84.34077 | 115.5959 | -1.33815 | 6.832903 | 8.63E-05 | 0.012927 |
| Myh14 | 65.3461 | 72.36009 | 45.87642 | 178.5313 | 137.4888 | 142.8451 | 97.54748 | 137.7358 | -1.1811 | 6.779772 | 0.000478 | 0.04545 |
| Car14 | 62.72621 | 54.48356 | 22.68834 | 239.2092 | 163.5859 | 105.3853 | 65.39598 | 91.64648 | -1.51027 | 6.655837 | 1.07E-05 | 0.002433 |
| Ranbp9 | 50.98709 | 57.95639 | 47.8754 | 85.74052 | 78.48951 | 340.0306 | 57.56303 | 52.15653 | -1.2316 | 6.592161 | 0.000282 | 0.031597 |
| Gadd45a | 35.92272 | 35.4684 | 85.85588 | 69.04728 | 116.1486 | 308.9597 | 36.8877 | 77.4897 | -1.21817 | 6.582497 | 0.000335 | 0.035293 |
| D230025D | 37.53496 | 60.34752 | 24.08762 | 41.07358 | 77.96096 | 405.8039 | 48.45496 | 38.95774 | -1.58806 | 6.522046 | 4.02E-06 | 0.001066 |
| Atp9a | 51.94436 | 51.58005 | 30.08454 | 186.4913 | 129.8248 | 98.72723 | 79.05809 | 95.90415 | -1.40434 | 6.501857 | 3.94E-05 | 0.006732 |
| Casq2 | 41.91824 | 38.99816 | 39.37976 | 126.4957 | 173.8265 | 72.02783 | 78.87593 | 106.6548 | -1.47636 | 6.408094 | 1.69E-05 | 0.003555 |
| Chrna1 | 17.23081 | 17.13642 | 18.19065 | 205.8682 | 135.1103 | 71.42256 | 71.95379 | 88.55966 | -2.70996 | 6.292174 | 1.54E-13 | 2.08E-10 |
| Myom3 | 32.39595 | 36.60703 | 15.59199 | 145.5087 | 131.4765 | 67.92541 | 49.63901 | 43.90728 | -1.63359 | 6.034742 | 2.28E-06 | 0.000678 |
| Spsb1 | 52.95201 | 21.80481 | 27.58582 | 63.49802 | 69.70239 | 179.0944 | 41.44174 | 49.33582 | -1.23876 | 5.984581 | 0.000266 | 0.030351 |
| Plxnd1 | 36.22502 | 44.34973 | 20.08968 | 119.4454 | 120.1788 | 56.35791 | 60.20438 | 37.94654 | -1.22969 | 5.954474 | 0.000291 | 0.032316 |
| Plbd2 | 37.78687 | 43.32496 | 19.19014 | 99.43172 | 69.76845 | 98.45821 | 57.83628 | 63.49259 | -1.21569 | 5.93836 | 0.000339 | 0.035329 |
| Kctd20 | 36.37616 | 50.72608 | 7.795994 | 41.43746 | 81.6608 | 152.5295 | 62.02599 | 36.50957 | -1.23619 | 5.876932 | 0.000265 | 0.030351 |
| Slc43a2 | 20.30415 | 22.37413 | 13.79291 | 58.1762 | 53.77985 | 176.337 | 34.0642 | 25.01385 | -1.8803 | 5.66163 | 8.62E-08 | 4.38E-05 |
| Slc47a1 | 29.12108 | 29.49058 | 17.39106 | 94.10989 | 69.0417 | 42.30206 | 45.08497 | 64.23769 | -1.31099 | 5.615314 | 0.000122 | 0.017038 |
| Peg3 | 22.62174 | 22.77265 | 3.6981 | 35.47884 | 73.53437 | 118.2306 | 27.41531 | 12.29404 | -1.69452 | 5.309186 | 1.07E-06 | 0.000385 |
| Ampd3 | 21.91639 | 15.31461 | 12.99332 | 47.30512 | 40.69827 | 103.0987 | 32.33367 | 25.27995 | -1.56719 | 5.229128 | 6.23E-06 | 0.001536 |
| Arl4d | 16.17278 | 12.01257 | 21.9887 | 33.61392 | 24.18111 | 151.857 | 11.8405 | 17.72258 | -1.5201 | 5.181967 | 1.21E-05 | 0.002655 |
| Sln | 4.332895 | 5.863957 | 10.59456 | 70.41185 | 106.3705 | 22.1262 | 18.21615 | 26.34437 | -2.82962 | 5.053165 | 5E-14 | 8.14E-11 |
| Nqo1 | 12.99868 | 13.09427 | 14.09276 | 87.96932 | 56.02618 | 25.48884 | 18.39831 | 24.10909 | -1.66487 | 4.986871 | 2.01E-06 | 0.000637 |
| Myh8 | 9.522292 | 12.41109 | 3.098408 | 48.21483 | 41.22681 | 19.57059 | 15.75697 | 68.92113 | -2.20083 | 4.783658 | 8.87E-10 | 6.77E-07 |
| Prepl | 11.94065 | 14.06211 | 2.89851 | 59.04042 | 53.11916 | 25.6906 | 19.58236 | 16.55172 | -1.83821 | 4.674952 | 1.79E-07 | 8.57E-05 |
| Fam174b | 2.771037 | 3.814419 | 2.798562 | 85.14921 | 47.30513 | 16.94773 | 10.83861 | 7.717038 | -3.42046 | 4.484424 | 2.29E-18 | 6.98E-15 |
| Ift122 | 10.32841 | 5.294641 | 2.698613 | 59.1314 | 33.16644 | 12.37453 | 19.03588 | 26.02505 | -2.27432 | 4.406394 | 3.34E-10 | 2.91E-07 |
| Ttc9 | 8.46426 | 9.678376 | 8.09584 | 36.52501 | 37.06449 | 15.1319 | 19.30912 | 20.91583 | -1.55913 | 4.289486 | 8.99E-06 | 0.002112 |
| Postn | 8.161965 | 5.807026 | 13.69296 | 24.83519 | 36.46987 | 21.52092 | 18.12507 | 26.07827 | -1.47353 | 4.283037 | 2.78E-05 | 0.005174 |
| Rab15 | 4.282512 | 8.198154 | 3.598151 | 45.4402 | 38.98048 | 23.2695 | 14.48184 | 16.01951 | -2.35661 | 4.282544 | 1.24E-10 | 1.16E-07 |
| Ifi30 | 6.549725 | 8.42588 | 11.39414 | 30.47541 | 19.55631 | 46.33723 | 10.56537 | 14.4761 | -1.47461 | 4.218663 | 2.71E-05 | 0.005125 |
| Myl6b | 6.801637 | 5.920889 | 7.29625 | 25.4265 | 27.88095 | 23.20224 | 26.32234 | 23.73654 | -1.92476 | 4.205808 | 8.11E-08 | 4.21E-05 |
| Myog | 6.499342 | 9.052128 | 5.597124 | 78.73573 | 21.73657 | 7.532322 | 5.009441 | 10.43131 | -1.80892 | 4.195008 | 3.66E-07 | 0.000157 |
| Pde8a | 12.19256 | 13.37893 | 2.19887 | 25.65393 | 27.1542 | 20.37762 | 19.03588 | 14.63576 | -1.18925 | 4.086525 | 0.000504 | 0.047607 |
| Tmem132a | 6.297812 | 6.148615 | 4.39774 | 37.38924 | 26.55958 | 7.46507 | 9.016994 | 11.22962 | -1.70257 | 3.781695 | 1.77E-06 | 0.000577 |
| Cables1 | 5.290162 | 6.547137 | 4.897483 | 16.23839 | 8.324645 | 45.32844 | 7.832944 | 7.025166 | -1.59867 | 3.680984 | 6.82E-06 | 0.001649 |
| Zfp52 | 5.139015 | 6.262479 | 6.59661 | 22.2425 | 21.20802 | 20.04136 | 9.654559 | 7.770259 | -1.43642 | 3.645144 | 5.13E-05 | 0.008293 |
| Ehd3 | 7.708522 | 7.059521 | 3.398254 | 23.78902 | 17.37605 | 13.9886 | 13.11563 | 8.462131 | -1.32704 | 3.587655 | 0.000145 | 0.019414 |
| Garem | 6.297812 | 6.945658 | 2.89851 | 9.91588 | 15.19578 | 28.98599 | 15.39265 | 7.82348 | -1.50167 | 3.560915 | 1.96E-05 | 0.003984 |
| Slc1a1 | 7.103932 | 4.668393 | 3.298305 | 15.46513 | 16.71536 | 14.25761 | 7.468621 | 8.728236 | -1.30421 | 3.303471 | 0.000214 | 0.026214 |
| Rian | 6.751255 | 4.839188 | 2.798562 | 18.7401 | 15.59219 | 12.77805 | 8.106187 | 6.546177 | -1.34794 | 3.275662 | 0.000129 | 0.017762 |
| Kcnk5 | 1.61224 | 1.76488 | 0.999486 | 3.638855 | 9.315674 | 38.93942 | 9.74564 | 4.151234 | -3.13156 | 3.149437 | 3.81E-15 | 8.47E-12 |
| Lpin2 | 4.23213 | 5.123846 | 3.798048 | 7.141253 | 9.183537 | 27.23742 | 7.468621 | 4.523781 | -1.3322 | 3.12251 | 0.000178 | 0.022856 |

**Down-regulated**

| Gene Name | WT1 | WT2 | WT3 | KO1 | KO2 | KO3 | KO4 | KO5 | logFC | logCPM | PValue | FDR |
|---|---|---|---|---|---|---|---|---|---|---|---|---|
| Tnnc2 | 17656.95 | 12717.16 | 38393.47 | 9056.747 | 8045.175 | 6075.222 | 9304.172 | 18551.55 | 1.167257 | 13.87028 | 0.000187 | 0.023598 |
| Pvalb | 10044.51 | 7370.539 | 22859.35 | 3678.109 | 3125.442 | 3081.998 | 3970.574 | 9689.087 | 1.511372 | 12.96172 | 1.33E-06 | 0.000457 |
| Myl1 | 9201.154 | 7349.872 | 7895.842 | 2179.31 | 2589.031 | 1996.267 | 3912.92 | 4232.609 | 1.450329 | 12.26436 | 3.47E-06 | 0.000953 |
| Myh1 | 5221.793 | 4463.041 | 2612.657 | 518.7643 | 1398.739 | 783.0253 | 3504.787 | 2596.331 | 1.219561 | 11.36496 | 9.52E-05 | 0.014092 |
| Car3 | 3529.344 | 2547.064 | 4079.404 | 396.6352 | 755.5607 | 388.3854 | 1943.936 | 1400.935 | 1.792784 | 10.87669 | 1.03E-08 | 6.61E-06 |
| Trdn | 2009.103 | 1694.57 | 5224.715 | 314.579 | 762.1675 | 805.4877 | 1236.877 | 1797.005 | 1.597808 | 10.75704 | 3.23E-07 | 0.000144 |
| Atp5e | 755.3848 | 628.7528 | 3112.5 | 616.2856 | 442.8579 | 344.2002 | 393.9242 | 1575.5 | 1.151575 | 9.942092 | 0.00023 | 0.027614 |
| Ppp1r3c | 1663.681 | 874.5267 | 891.4419 | 668.2303 | 506.6141 | 109.3532 | 293.28 | 470.6329 | 1.480598 | 9.419704 | 2.19E-06 | 0.000674 |

| | | | | | | | | | | | | |
|---|---|---|---|---|---|---|---|---|---|---|---|---|
| Rps21 | 304.008 | 234.2167 | 1166.401 | 254.7199 | 182.3494 | 182.7933 | 194.1842 | 492.6663 | 1.1198 | 8.556313 | 0.000343 | 0.035668 |
| Asb5 | 501.8601 | 465.9284 | 667.9567 | 15.23771 | 282.5755 | 198.127 | 400.1177 | 353.3339 | 1.125926 | 8.49469 | 0.00032 | 0.034161 |
| Phka1 | 560.4045 | 655.5676 | 461.6628 | 124.9037 | 236.856 | 194.0246 | 332.1715 | 251.6287 | 1.295191 | 8.460517 | 3.47E-05 | 0.006118 |
| Ndufa2 | 243.1459 | 205.5801 | 1097.936 | 219.8778 | 142.0475 | 134.4385 | 147.733 | 500.1705 | 1.170821 | 8.394009 | 0.000181 | 0.023073 |
| Rpl37a | 222.741 | 176.4311 | 1052.159 | 222.3795 | 164.1144 | 156.2957 | 159.4824 | 434.8684 | 1.088061 | 8.33799 | 0.000505 | 0.047607 |
| Dhrs7c | 289.4475 | 249.5313 | 321.7347 | 115.4882 | 122.6233 | 42.50382 | 141.2662 | 206.1247 | 1.19147 | 7.540687 | 0.000141 | 0.019182 |
| Smox | 280.0259 | 295.3043 | 156.8194 | 120.9464 | 100.9528 | 94.15403 | 121.0463 | 119.8004 | 1.131959 | 7.333422 | 0.000302 | 0.032912 |
| Gpd2 | 185.9618 | 185.9387 | 162.5165 | 36.61598 | 56.22439 | 79.08939 | 128.0595 | 104.5792 | 1.139437 | 6.876216 | 0.00028 | 0.031597 |
| Myeov2 | 96.28096 | 65.75602 | 378.5055 | 65.49939 | 57.6779 | 47.48053 | 50.18549 | 167.646 | 1.211124 | 6.859926 | 0.000111 | 0.015894 |
| Aldh1a1 | 179.2106 | 198.0082 | 140.3279 | 35.25141 | 53.71378 | 50.64142 | 73.04676 | 93.77532 | 1.493565 | 6.688209 | 1.93E-06 | 0.000619 |
| Mrln | 84.94489 | 84.82812 | 237.378 | 8.096453 | 14.7333 | 9.617162 | 46.17794 | 96.59603 | 1.951817 | 6.186224 | 5.86E-10 | 4.61E-07 |
| Sorbs2 | 114.6202 | 132.2522 | 90.75336 | 22.60639 | 42.87853 | 24.95082 | 82.337 | 68.92113 | 1.220902 | 6.180433 | 0.000103 | 0.014844 |
| Tspan8 | 82.07309 | 67.57784 | 145.8251 | 4.775997 | 23.25615 | 17.21674 | 59.93113 | 64.82312 | 1.535296 | 5.863642 | 1.09E-06 | 0.000385 |
| Crip1 | 37.88764 | 34.84215 | 163.7159 | 33.3865 | 29.73088 | 19.8396 | 30.42097 | 58.59627 | 1.191774 | 5.674546 | 0.000151 | 0.020007 |
| Ankrd1 | 29.67529 | 25.67616 | 188.1033 | 21.6057 | 66.13468 | 16.4097 | 19.94668 | 39.70283 | 1.304037 | 5.669479 | 3.39E-05 | 0.006039 |
| Nrep | 116.2324 | 87.84549 | 85.85588 | 19.78627 | 19.68845 | 8.137598 | 30.69421 | 32.94377 | 2.119466 | 5.651579 | 2.11E-11 | 2.31E-08 |
| Mettl21c | 81.72041 | 99.7442 | 108.7441 | 2.63817 | 10.1085 | 1.950333 | 27.77963 | 44.54594 | 2.475654 | 5.561824 | 7.7E-15 | 1.45E-11 |
| 2210408F2 | 37.9884 | 39.33975 | 102.3474 | 25.29004 | 24.18111 | 18.89806 | 23.68099 | 45.92968 | 1.113665 | 5.313466 | 0.000409 | 0.04111 |
| Asb15 | 73.20577 | 63.53569 | 47.57555 | 4.821483 | 20.67948 | 12.10552 | 46.72442 | 36.56279 | 1.350156 | 5.257781 | 1.96E-05 | 0.003984 |
| 1500012F0 | 24.13322 | 21.69095 | 117.0399 | 17.1481 | 16.8475 | 34.50073 | 13.66211 | 33.10343 | 1.229075 | 5.120116 | 9.76E-05 | 0.014269 |
| Nrtn | 28.56688 | 25.3915 | 103.9466 | 12.28114 | 9.183537 | 10.02068 | 13.66211 | 41.88489 | 1.58849 | 4.937358 | 4.97E-07 | 0.000192 |
| G0s2 | 54.21157 | 22.26026 | 66.166 | 22.28799 | 15.59219 | 6.523529 | 18.21615 | 31.40036 | 1.333952 | 4.89112 | 2.42E-05 | 0.004682 |
| Flrt1 | 60.35823 | 42.98338 | 69.96405 | 1.773942 | 2.180264 | 2.219345 | 21.58614 | 20.11752 | 2.601069 | 4.792533 | 7.9E-16 | 1.93E-12 |
| Amy1 | 42.42206 | 36.77783 | 57.37052 | 6.277025 | 8.919263 | 15.80443 | 24.04532 | 28.84576 | 1.440848 | 4.788353 | 5.52E-06 | 0.001434 |
| Mss51 | 60.25747 | 46.34234 | 66.76569 | 0.409371 | 0.726755 | 1.008793 | 11.47617 | 6.492956 | 3.878137 | 4.600081 | 2.68E-30 | 3.27E-26 |
| Retnla | 18.38961 | 18.61664 | 84.6565 | 10.18879 | 16.05467 | 7.46507 | 12.7513 | 24.58808 | 1.504704 | 4.590681 | 2.08E-06 | 0.000651 |
| 2310065F0 | 13.04907 | 9.849171 | 40.0794 | 7.687081 | 5.21942 | 5.38023 | 6.193491 | 16.44527 | 3.159559 | 4.553124 | 8.09E-22 | 4.94E-18 |
| Kdm1a | 38.49223 | 35.86692 | 42.47817 | 12.82696 | 12.68517 | 16.74597 | 14.02644 | 11.12318 | 1.530425 | 4.531719 | 1.45E-06 | 0.000493 |
| Gm15417 | 24.23398 | 23.51276 | 66.166 | 8.960681 | 8.126439 | 8.742874 | 12.84239 | 23.6301 | 1.601031 | 4.463716 | 4.66E-07 | 0.000187 |
| Arhgap26 | 58.29255 | 24.4806 | 18.99024 | 10.09782 | 17.77246 | 12.8453 | 17.12318 | 9.739434 | 1.333993 | 4.413969 | 2.82E-05 | 0.005215 |
| Kcnf1 | 98.64893 | 28.12422 | 9.994864 | 2.00137 | 4.162323 | 0.538023 | 11.74942 | 5.854305 | 3.256271 | 4.346476 | 1.07E-22 | 8.71E-19 |
| 9530091C0 | 34.05857 | 31.76785 | 21.38901 | 8.278395 | 11.29773 | 13.58508 | 9.563479 | 10.96352 | 1.439054 | 4.149796 | 6.59E-06 | 0.001609 |
| Sh2d7 | 18.28885 | 15.77006 | 45.07684 | 12.09919 | 9.447812 | 7.061552 | 10.92969 | 21.39482 | 1.104255 | 4.13525 | 0.000522 | 0.048444 |
| 2310009B1 | 54.06042 | 51.40926 | 51.97329 | 2.183313 | 4.624803 | 1.076046 | 11.93158 | 9.792655 | 1.34162 | 3.703934 | 2.85E-05 | 0.005229 |
| Lrrc38 | 19.49803 | 22.54492 | 13.99281 | 11.37142 | 8.390714 | 3.160885 | 11.11185 | 8.834678 | 1.125316 | 3.643881 | 0.000469 | 0.04545 |
| Zfp423 | 26.95464 | 16.85176 | 9.994864 | 5.185369 | 8.456782 | 6.052759 | 12.56914 | 8.994341 | 1.132028 | 3.587767 | 0.000476 | 0.04545 |
| E030003E1 | 24.28436 | 26.24548 | 1.49923 | 3.729827 | 6.474724 | 6.658035 | 9.199156 | 8.621794 | 1.338834 | 3.462604 | 3.88E-05 | 0.006727 |
| Pf4 | 7.557375 | 9.963034 | 29.28495 | 5.731197 | 5.615832 | 4.035173 | 7.104298 | 11.81505 | 1.16899 | 3.347667 | 0.000292 | 0.032316 |
| Cilp2 | 11.48721 | 12.52496 | 18.39055 | 6.367997 | 4.75694 | 4.102426 | 9.563479 | 6.971945 | 1.152492 | 3.225082 | 0.000403 | 0.040708 |
| Polk | 17.33158 | 14.51756 | 9.595069 | 3.775312 | 6.21045 | 3.093632 | 9.108075 | 5.215653 | 1.349295 | 3.126182 | 3.85E-05 | 0.006715 |
| Npnt | 13.70404 | 17.6488 | 7.196302 | 6.004111 | 7.069342 | 4.304184 | 6.375652 | 5.109211 | 1.162053 | 3.100161 | 0.00038 | 0.038956 |
| 1600002K0 | 8.46426 | 5.579299 | 24.88721 | 4.275655 | 4.030185 | 4.976713 | 5.738087 | 9.52655 | 1.165833 | 3.08322 | 0.000334 | 0.035288 |

**Table S2. Cluster 6 genes in Figure 5A**

| Gene Name | WTex_1 | WTex_2 | WTex_3 | KOex_1 | KOex_2 | KOex_3 |
|---|---|---|---|---|---|---|
| Acadsb | 88.05 | 91.56 | 94.38 | 108.89 | 103.39 | 115 |
| Acsl1 | 408.16 | 414.23 | 373.86 | 477.07 | 411.83 | 494.87 |
| Actn3 | 55.62 | 44.73 | 131.93 | 322.82 | 107.74 | 224.24 |
| Adk | 121.63 | 137.51 | 115.77 | 128.39 | 124.97 | 139.36 |
| Aes | 948.41 | 963.85 | 891.82 | 1012.32 | 1085.93 | 1098.6 |
| Akap1 | 75.47 | 89.73 | 69.13 | 100.45 | 92.24 | 130.58 |
| Amotl1 | 161.68 | 162.5 | 146.3 | 191.03 | 160.44 | 218.09 |
| Ankrd23 | 1971.41 | 1989.87 | 1368.67 | 2157.12 | 2350.68 | 2188.78 |
| Ankrd40 | 158.83 | 161.21 | 150.14 | 166.38 | 175.49 | 190.79 |
| Arl1 | 126.49 | 153.67 | 138.79 | 137.38 | 157.03 | 145.97 |
| Asb11 | 208.81 | 162.19 | 173.67 | 227.74 | 209.39 | 199.57 |
| Asb5 | 222.03 | 183.92 | 146.81 | 331.43 | 193.96 | 196.08 |
| Atcayos | 145.76 | 121.4 | 97.01 | 156.28 | 162.9 | 117.24 |
| Atp5c1 | 1501.11 | 1412.22 | 1422.63 | 1531.89 | 1554.23 | 1624.41 |
| Atp5k | 1835.47 | 1831.35 | 1990.74 | 1884.12 | 1974.69 | 2012.15 |
| Bdh1 | 196.48 | 152.56 | 146.57 | 204.23 | 181.68 | 250.12 |
| Bnip3 | 765.56 | 632.28 | 697.1 | 867.49 | 808.79 | 879.57 |
| Btbd1 | 266.42 | 252.07 | 247.43 | 288.01 | 289.05 | 311.15 |
| Bzw2 | 161 | 175.15 | 159.27 | 202.03 | 196.91 | 192.86 |
| Cap2 | 176.5 | 180.55 | 172.85 | 189.84 | 210.93 | 214.81 |
| Cat | 223.31 | 237.43 | 222.35 | 230.75 | 265.85 | 270.3 |
| Cd24a | 51.04 | 47.74 | 57.06 | 85.37 | 80 | 83.12 |
| Cd36 | 593.92 | 586.69 | 578.15 | 635.83 | 666.94 | 789.91 |
| Chchd3 | 505.5 | 467.89 | 449.75 | 522.88 | 505.27 | 556.84 |
| Cisd3 | 101.78 | 112.63 | 101.98 | 115.49 | 118.5 | 126.36 |
| Clu | 45.59 | 54.27 | 38.6 | 58.42 | 77.37 | 75.86 |
| Cox19 | 112.42 | 115.48 | 121.46 | 148.87 | 135.4 | 142.38 |
| Cox7a2 | 505.34 | 538.25 | 533.21 | 526.51 | 558.15 | 551.07 |
| Cox7b | 910.55 | 931.48 | 978.17 | 964.7 | 1043.42 | 985.04 |
| Cpe | 135.9 | 153.7 | 136.58 | 179.46 | 183.48 | 206.77 |
| Ctdnep1 | 187.61 | 220.01 | 197.43 | 205.32 | 220.69 | 212.43 |
| Cul3 | 112.58 | 108.32 | 109.43 | 121.95 | 130.49 | 130.05 |
| Cycs | 541.12 | 505.27 | 503.93 | 537.32 | 517.87 | 567.68 |

| | | | | | | |
|---|---|---|---|---|---|---|
| Cyfip2 | 17.36 | 19.71 | 42.3 | 79.22 | 41.65 | 42.02 |
| Dazap2 | 162.69 | 204.43 | 179.93 | 192.94 | 200.14 | 206.05 |
| Ddit4l | 26.84 | 41.07 | 21.57 | 32.2 | 63.9 | 43.32 |
| Dmpk | 318.17 | 332.23 | 331.53 | 389.67 | 399.03 | 387.3 |
| Dnaja4 | 210.57 | 203.27 | 195.63 | 235.73 | 226.62 | 242.52 |
| Dusp13 | 53.86 | 64.36 | 56.9 | 74.77 | 75.88 | 65.63 |
| Dusp18 | 58.96 | 33.58 | 30.57 | 71.32 | 45.36 | 67.46 |
| Dusp26 | 34.37 | 32.17 | 36.33 | 70.64 | 53.27 | 48.78 |
| E2f6 | 154.39 | 144 | 130.82 | 153.74 | 176.02 | 192.87 |
| Eif4g2 | 211.6 | 228.4 | 215.48 | 220.59 | 231.55 | 250.3 |
| Entpd4 | 66.58 | 102.61 | 103.82 | 86.46 | 75.5 | 133.21 |
| Esr1 | 6.6 | 7.86 | 5.69 | 89.62 | 138.11 | 128.69 |
| Esrrg | 47.77 | 47.44 | 41.54 | 93.31 | 73.14 | 99.77 |
| Fam134b | 396.85 | 352.1 | 319.25 | 444.66 | 376.42 | 397.57 |
| Fam213a | 82.81 | 65.06 | 69.08 | 87.15 | 97.71 | 94.03 |
| Fam220a | 94.49 | 95.94 | 77.65 | 120.6 | 124.71 | 129.08 |
| Fermt2 | 142.28 | 151.95 | 146.16 | 160.11 | 163.57 | 165.26 |
| Fkbp3 | 518.51 | 556.17 | 536.5 | 557.88 | 643.45 | 586.96 |
| Fth1 | 3581.78 | 3674.52 | 3562.46 | 3970.07 | 3977.59 | 3935.87 |
| Fxyd6 | 306.09 | 297.43 | 253.54 | 397.02 | 347.88 | 376.6 |
| Fzd9 | 61.22 | 64.08 | 47.19 | 104.61 | 79.96 | 96.4 |
| Gabarap | 332.93 | 363.48 | 339.56 | 353.69 | 365.78 | 347.66 |
| Gabarapl1 | 187.45 | 140.15 | 106.04 | 189.44 | 159.03 | 173.08 |
| Gapdh | 2072.01 | 1950.17 | 2015.55 | 2179.76 | 2233.59 | 2226.01 |
| Gng5 | 526.32 | 624.33 | 554.7 | 583.05 | 623.11 | 658.71 |
| Got2 | 887.85 | 853.51 | 841.5 | 962.18 | 937.35 | 1013.13 |
| Grsf1 | 142.67 | 133.96 | 133.97 | 146.57 | 157.19 | 166.6 |
| Gstm1 | 141.22 | 151.21 | 143.43 | 173.91 | 189.69 | 166.67 |
| Gstm5 | 95.83 | 95.51 | 104.95 | 120.27 | 116.71 | 124.71 |
| Gyg | 654.79 | 604.58 | 613.55 | 652.45 | 686.44 | 674.04 |
| Hba-a1 | 47.59 | 23.5 | 0.01 | 9.66 | 296.11 | 771.75 |
| Hint1 | 361.76 | 338.16 | 379.29 | 401.62 | 392.96 | 367.46 |
| Hrc | 700.95 | 651.68 | 624.85 | 715.93 | 714.05 | 794.52 |
| Hspa8 | 1367.82 | 1356.61 | 1369.23 | 1404.3 | 1745.68 | 1682.53 |
| Hspb6 | 3581.59 | 3510.66 | 3619.27 | 4146.72 | 4102.84 | 4061.11 |
| Hspb8 | 304.11 | 333.44 | 322.3 | 350.14 | 367.17 | 350.16 |

| | | | | | | |
|---|---|---|---|---|---|---|
| Idh2 | 1504.72 | 1653.9 | 1562.26 | 1598.56 | 1979.11 | 1777.74 |
| Idh3a | 416.03 | 410.44 | 389.37 | 445 | 448.26 | 541.38 |
| Kif1b | 145.53 | 180.47 | 159.99 | 182.21 | 174.73 | 193.81 |
| Ldb3 | 1752.67 | 1825.75 | 2002.14 | 1846.47 | 1879.71 | 1983.64 |
| Lynx1 | 198.27 | 184.41 | 145.45 | 213.3 | 207.03 | 248.33 |
| Map1lc3a | 399.95 | 359.82 | 380.32 | 451.16 | 468.9 | 446.17 |
| Mgll | 67.35 | 84.06 | 66 | 85.01 | 66.07 | 81.91 |
| Mif | 173.77 | 196.15 | 171.1 | 210.86 | 218.54 | 230.37 |
| Mir2861 | 0 | 0 | 0 | 17.61 | 17.36 | 22.28 |
| Mir8094 | 0 | 0 | 19.76 | 7.85 | 8.9 | 18.78 |
| Morf4l1 | 267.96 | 292.85 | 283.18 | 299.55 | 321.67 | 306.91 |
| Mpc1 | 859.11 | 889.15 | 928.04 | 922.21 | 1023.66 | 971.84 |
| Mrpl30 | 305.62 | 322.48 | 293.98 | 317.3 | 302.83 | 319.59 |
| Mrpl42 | 544.04 | 613.19 | 566.66 | 647.04 | 591.41 | 634.48 |
| Mtch2 | 141.52 | 136.06 | 129.02 | 154.46 | 144.07 | 154.03 |
| Murc | 118.4 | 131.09 | 116.8 | 173.18 | 149.45 | 152.59 |
| Myot | 2023.25 | 1950.58 | 2146.12 | 2255.11 | 2224.66 | 2366.6 |
| Naa50 | 153.98 | 139.1 | 124.23 | 158.62 | 168.31 | 185.3 |
| Ndrg4 | 23.36 | 29.57 | 32.82 | 38.16 | 73.29 | 50.48 |
| Ndufa3 | 959.74 | 939.42 | 886.09 | 972.47 | 991.22 | 975.6 |
| Ndufa5 | 775.49 | 820.27 | 750.87 | 809.66 | 818.56 | 801.37 |
| Ndufa6 | 896.16 | 899.66 | 879.54 | 915.89 | 933.58 | 899.82 |
| Ndufb4 | 739.73 | 794.13 | 792.74 | 820.87 | 833.92 | 829.14 |
| Ndufv2 | 562.94 | 566.81 | 588.23 | 615.74 | 614.9 | 638.66 |
| Neb | 540.27 | 463.38 | 493.38 | 531.55 | 551.54 | 698.56 |
| Nexn | 461.85 | 429.44 | 463.98 | 515.93 | 473.33 | 534.04 |
| Nnt | 417.12 | 427.67 | 434.16 | 471.59 | 448.62 | 456.13 |
| Nqo1 | 21.26 | 31.84 | 24.78 | 32.62 | 69.59 | 47.64 |
| Nrap | 1505.16 | 1307.6 | 1288.09 | 1497.6 | 1465.14 | 1423.95 |
| Nudt4 | 194.85 | 199.33 | 210.58 | 242.94 | 222.42 | 246.97 |
| Optn | 91.47 | 78.48 | 72.82 | 98.74 | 89.93 | 102.9 |
| Pcp4l1 | 222.17 | 211.72 | 223.64 | 306.94 | 331.97 | 320.03 |
| Pdzrn3 | 54.74 | 71.87 | 53.93 | 81.22 | 80.18 | 90.69 |
| Perm1 | 155.35 | 144.3 | 103.7 | 179.57 | 178.36 | 218.34 |
| Pfn2 | 175.15 | 180.79 | 203.46 | 193.57 | 213.25 | 194.02 |
| Pgk1 | 561.13 | 537.43 | 555.8 | 617.01 | 575.13 | 604.21 |

| | | | | | | |
|---|---|---|---|---|---|---|
| Phospho1 | 134.39 | 132.61 | 125.18 | 179.08 | 152.24 | 173.51 |
| Phyh | 389.13 | 352.53 | 338.02 | 403.77 | 459.18 | 516.64 |
| Pkia | 269.56 | 345.11 | 306.86 | 292.6 | 343.59 | 341.73 |
| Pla2g16 | 144.89 | 138.84 | 122.95 | 162.58 | 162.53 | 186.63 |
| Plekhb2 | 65.67 | 63.7 | 55.73 | 71.45 | 81.23 | 90.47 |
| Plin5 | 81.79 | 53.56 | 64.96 | 78.62 | 72.59 | 119.01 |
| Ppm1a | 182.18 | 171 | 165.33 | 180.32 | 206.52 | 201.71 |
| Ppm1b | 299.11 | 286.26 | 261.94 | 297.44 | 366.93 | 354.76 |
| Ppp1r14b | 86.52 | 117.09 | 90.86 | 129.5 | 112.91 | 104.74 |
| Ppp1r27 | 272.79 | 397.5 | 196.25 | 554.48 | 342 | 427.2 |
| Ppp2r3a | 160.01 | 168.66 | 160.32 | 180.96 | 173.27 | 197.52 |
| Pptc7 | 87.34 | 97.65 | 75.92 | 103.69 | 96.66 | 133.37 |
| Prdx3 | 292.37 | 291.99 | 270.48 | 293 | 311.85 | 339.7 |
| Psma1 | 171.62 | 172.21 | 186.12 | 188.26 | 191.23 | 177.57 |
| Psme1 | 94.61 | 112.33 | 100.14 | 110.4 | 119.53 | 139.58 |
| Qk | 160.53 | 173.5 | 174.19 | 185.71 | 161.74 | 204.83 |
| Rangrf | 81.29 | 84.51 | 69.71 | 114.96 | 104.89 | 106.87 |
| Rassf3 | 79.12 | 89.05 | 76.29 | 88.85 | 101.48 | 104.63 |
| Rcan1 | 282.18 | 163.78 | 120.56 | 320.83 | 205.93 | 247.6 |
| Reep5 | 80.73 | 94.88 | 91.59 | 109.12 | 113.68 | 121.85 |
| Rilpl1 | 169.36 | 145.28 | 121.26 | 184.92 | 186.33 | 182.61 |
| Rnf11 | 140.68 | 149.39 | 125.72 | 148.58 | 164.81 | 160.03 |
| Rnf115 | 109.65 | 106.07 | 118.09 | 168.72 | 113.34 | 132.4 |
| Rps13 | 756.22 | 712.55 | 780.43 | 740.54 | 824.38 | 763.52 |
| Rragd | 63.54 | 60.2 | 55.97 | 76.12 | 76.05 | 85.33 |
| Rwdd1 | 201.7 | 206.14 | 194.78 | 218.04 | 224.98 | 216.6 |
| S100a1 | 569.19 | 489.43 | 497.26 | 759.53 | 728.24 | 728.38 |
| Sdr39u1 | 123.43 | 122.59 | 110.48 | 125.12 | 140.52 | 149.78 |
| Sepw1 | 1371.11 | 1374.04 | 1283.37 | 1412.39 | 1483.5 | 1344.99 |
| Skp1a | 323.79 | 323.02 | 342.31 | 324.07 | 386.57 | 346.97 |
| Slc6a8 | 75.21 | 87.72 | 68.3 | 97.72 | 91.05 | 99.43 |
| Smyd1 | 313.04 | 351.75 | 369.79 | 368.65 | 372.52 | 394.63 |
| Snx3 | 141.36 | 164.01 | 146.71 | 154.35 | 169.13 | 174.82 |
| Sod1 | 548.26 | 535.21 | 530.46 | 556 | 576.29 | 541.57 |
| Sord | 87.02 | 100 | 88.47 | 110.46 | 107.86 | 120.67 |
| Srl | 597.5 | 568.03 | 567.31 | 625.04 | 576.17 | 624.18 |

| | | | | | | |
|---|---|---|---|---|---|---|
| Stau2 | 83.83 | 75.84 | 91.68 | 109.19 | 108.31 | 104.09 |
| Sucla2 | 536.24 | 521.44 | 513.9 | 560.7 | 611.95 | 618.91 |
| Tead1 | 65.91 | 76.36 | 70.37 | 78.49 | 86.7 | 95.15 |
| Timm17a | 342.05 | 334.33 | 323.59 | 366.01 | 323.88 | 350.11 |
| Tmem126a | 147.52 | 133.2 | 140.32 | 169.86 | 155.6 | 170.72 |
| Tmod1 | 104.07 | 92.61 | 106.81 | 139.64 | 108.09 | 135.91 |
| Tnnc1 | 6489.1 | 6619.09 | 6499.46 | 7071.47 | 8785.46 | 6105.01 |
| Tnnt1 | 6462.35 | 5685.51 | 6269.47 | 6925.19 | 7636.19 | 5854.22 |
| Tpm3 | 8520.09 | 7330.48 | 7872.43 | 8823.33 | 10660.61 | 8210.4 |
| Tpt1 | 4875.8 | 5144.89 | 5338.24 | 5319.9 | 5475.75 | 5462.82 |
| Tspan7 | 137.45 | 162.29 | 140.65 | 160.41 | 147.8 | 159 |
| Ttc9 | 48.41 | 47.45 | 60.66 | 80.57 | 66.88 | 62.29 |
| Ttn | 255.43 | 203.67 | 207.39 | 255.88 | 263.38 | 328.08 |
| Tuba8 | 257.82 | 225.62 | 197.53 | 316.69 | 220.85 | 280.32 |
| Tubb4b | 158.24 | 164.68 | 150.58 | 194.02 | 135.06 | 201.05 |
| Ubc | 860.36 | 906.22 | 863.44 | 950.09 | 1046.47 | 943.34 |
| Ube2d3 | 256.31 | 278.71 | 286.47 | 292.61 | 271.72 | 295.81 |
| Uchl1 | 120.38 | 109.64 | 145.73 | 173.85 | 158.45 | 137.36 |
| Ucp2 | 74.37 | 86.62 | 111.74 | 75.54 | 102.89 | 143.63 |
| Ugp2 | 691.48 | 699.18 | 699.24 | 807.78 | 838.8 | 969.11 |
| Uqcc2 | 434.63 | 450.11 | 426.34 | 460.67 | 440.02 | 466.26 |
| Uqcrb | 1694.44 | 1528.52 | 1679.2 | 1721.9 | 1742.56 | 1687.73 |
| Uqcrc2 | 835.87 | 826.96 | 847.24 | 831.07 | 860.5 | 922.53 |
| Vegfa | 87.18 | 104.84 | 82.26 | 118.99 | 81.97 | 105.76 |
| Wipi1 | 86.35 | 101.13 | 87.88 | 100.5 | 112.51 | 110.33 |
| Wnk1 | 65.31 | 68.47 | 63.34 | 79.94 | 82.41 | 91.94 |
| Yipf7 | 224.73 | 219.97 | 236.33 | 267.8 | 220.86 | 257.44 |

**Table S3. Primers used in this study**

| Gene Symbol | Gene Name | Forward (5'-3') | Reverse (5'-3') |
|---|---|---|---|
| 36B4 | ribosomal protein, large, P0 | GCGTCCTGGCATTGTCTGT | GCAAATGCAGATGGATCAGCC |
| Lsd1 | lysine (K)-specific demethylase 1A | ATGGATGTCACACTTCTGGA | CAAGACCTGTTACAACCATG |
| Atrogin1 | F-box protein 32 | AGTGAGGACCGGCTACTGTG | GATCAAACGCTTGCGAATCT |
| MuRF1 | tripartite motif-containing 63 | CCTGCAGAGTGACCAAGGA | GGCGTAGAGGGTGTCAAACT |
| SMART | F-box protein 21 | TCAATAACCTCAAGGCGTTC | GTTTTGCACACAAGCTCCA |
| Ddit4 | DNA damage inducible transcript 4 | CCAGAGAAGAGGGCCTTGA | CCATCCAGGTATGAGGAGTCTT |
| Klf15 | Kruppel like factor 15 | ACAGGCGAGAAGCCCTTT | CATCTGAGCGGGAAAACCT |
| Gadd45a | growth arrest and DNA damage inducible alpha | CGTAGACCCCGATAACGTGGTA | CGGATGAGGGTGAAATGGAT |
| Gabarapl1 | GABA type A receptor associated protein like 1 | CTTTCCCCTTGTTTACCCTCCAT | CCCAATGTCAACCCCTTC |
| Hspb7 | heat shock protein family, member 7 | TGTCACCACCTTCAACAACCAC | TCATGACTGTGCCATCAGCTG |
| Dusp4 | dual specificity phosphatase 4 | CATCTGCCTGCTTAAAGGTGG | CTCATTGGTGCTGGGAGGTAC |
| Ampd3 | adenosine monophosphate deaminase 3 | ACGCTTGCTGGTCGGTTTAG | TGGCTTCCTTCTGTCCGATG |
| Runx1 | runt related transcription factor 1 | ACGATGAAAACTACTCGGCAG | CTGAGGTCGTTGAATCTCGCT |
| Myh7 | myosin, heavy polypeptide 7, cardiac muscle, beta | CTCAAGCTGCTCAGCAATCTATTT | GGAGCGCAAGTTTGTCATAAGT |
| Myh2 | myosin, heavy polypeptide 2, skeletal muscle, adult | GAGCAAAGATGCAGGGAAAG | TAAGGGTTGACGGTGACACA |
| Myl2 | myosin, light polypeptide 2, regulatory, cardiac, slow | TGACCACACAAGCAGAGAGG | CCGTGGGTAATGATGTGGA |
| Tnnc1 | troponin C, cardiac/slow skeletal | CTGCAGGAGATGATTGACGA | CAATGTAGCCATCAGCGTTT |
| Tnni1 | troponin I, skeletal, slow 1 | TGAAGCCAAATGCCTCCACAACAC | ACACCTTGTGCTTAGAGCCCAGTA |
| Sln | Sarcolipin | GCTCCTCTTCAGGAAGTGAAG | TGGCCCCTCAGTATTGGTAGG |
| Myh1 | myosin, heavy polypeptide 1, skeletal muscle, adult | GCATCGAGTGGGAGTTCATT | TGAGCTTCAACCTTGCCTTT |
| Myh4 | myosin, heavy polypeptide 4, skeletal muscle | ACATTATTGGCTGGCTGGAC | ACCCTTCTTCTTGCCACCTT |
| Myl1 | myosin, light polypeptide 1 | CCCAGCCAAACCTAAGGAAG | TGCATTCACCTGTTCTGTCAA |
| Mylpf | myosin light chain, phosphorylatable, fast skeletal muscle | AGGGAGCTCCAACGTCTTCT | GCGTCGAGTTCCTCATTCTT |
| Tnnc2 | troponin C2, fast | AGCGGAGGAGTCCCAGTC | ATCACGGTGCCCAACTCTT |
| Tnni2 | troponin I, skeletal, fast 2 | AGCAGCAAGGAGCTGGAAGA | ATGGCGTCGGCAGACATAC |
| Foxk1 | forkhead box K1 | ACCCACGAATAGCTTGACTG | GCATTAGCGGCTACTGAGACG |
| Nfatc1 | nuclear factor of activated T cells,cytoplasmic, calcineurin dependent 1 | GCCTTTTGCGAGCAGTATCT | CATAGTGAGCCCTGTGGTGA |
| Pitx1 | paired-like homeodomain transcription factor 1 | TACGAGGACGTGTACGCT | TGAAGAAGGTAAAGCTCTTGGT |
| Esrrg | estrogen-related receptor gamma | TGACTTGGCTGACCGAG | CCGAGGATCAGAATCTCC |
| Esrra | estrogen related receptor, alpha | CCTTCTTCAAGAGGACCATCC | TGGTGATCTCACACTCATTGG |
| Pgc-1a | peroxisome proliferative activated receptor, gamma, coactivator 1 alpha | AAGTGTGGAACTCTCTGGAACTG | GGGTTATCTTGGTTGGCTTTATG |
| Dhrs7c | dehydrogenase/reductase (SDR family) member 7C | CCCCTGAGCAAAGAAACTGG | CTTCTCCTTCACCCCACAGG |
| Smox | spermine oxidase | CAGTTCACAGGGAACCCCAAT | AGGAATAGGAACCCCGGAAGTA |
| Ankrd1 | ankyrin repeat domain 1 (cardiac muscle) | ACAGCGCGTGTCCTTTTGAT | GCCAAGGAAAACTGAGTTCGA |
| Asb15 | ankyrin repeat and SOCS box-containing 15 | ATCGTCCGGCTGCTTCTCT | GCAGCCTCAGCATGATCTCA |
| Ndufa6 | NADH:ubiquinone oxidoreductase subunit A6 | GTCACAGACCCCAGAGTGGT | TAACATGCACCTTCCCATCA |
| Cox19 | cytochrome c oxidase assembly protein 19 | AGCTTCCCGCTGGACCACTTC | CCTTTGCATCCTGCACATTAAATAC |
| Idh2 | isocitrate dehydrogenase 2 (NADP+), mitochondrial | GGAGAAGCCGGTAGTGGAGAT | GGTCTGGTCACGGTTTGGAA |
| Idh3a | isocitrate dehydrogenase 3 (NAD+) alpha | CGCGTGGGTGTCCAAGGTCTC | TGTGACATTGCGCTCCTCCAA |
| Ldhb | lactate dehydrogenase B | TCTTCCAACACATCCACCAG | GTCCCGAACAACAAGATCACT |
| Perm1 | PPARGC1 and ESRR induced regulator, muscle 1 | GAAAACAGTCTTGCTCGCCAA | CTTGTCACCTGGGTTCGTCT |
| Vegfa | vascular endothelial growth factor A | GGCAGCTTGAGTTAAACGAAC | TGGTGACATGGTTAATCGGTC |

**Table S4. ChIP primers used in this study**

| Gene Name | Forward (5'-3') | Reverse (5'-3') |
|---|---|---|
| Sln | ATTGGCAGCCAAAGGAGCAG | TCTCAAAGGGACAAGCTCCA |
| Gadd45a | ACTACCTAGAGCTGGCTCAG | CAAGGCAGAGTTGAGCGGA |
| Ddit4 | CCACACCCCCTTTCTGGAA | AAGTCTTTCCTTGATCCTCCC |
